# GSAP regulates mitochondrial function through the Mitochondria-associated ER membrane in the pathogenesis of Alzheimer’s disease

**DOI:** 10.1101/2020.11.17.385245

**Authors:** Peng Xu, Jerry C. Chang, Xiaopu Zhou, Wei Wang, Michael Bamkole, Eitan Wong, Karima Bettayeb, Lu-Lin Jiang, Timothy Huang, Wenjie Luo, Huaxi Xu, Angus C. Nairn, Marc Flajolet, Nancy Ip, Yue-Ming Li, Paul Greengard

## Abstract

Biochemical, pathogenic and human genetic data confirm that GSAP (γ-secretase activating protein), a selective γ-secretase modulatory protein, plays important roles in Alzheimer’s disease (AD) and Down syndrome. However, the molecular mechanism(s) underlying GSAP-dependent pathogenesis remains largely elusive. Here, through unbiased proteomics and single-nuclei RNA-seq, we identified that GSAP regulates multiple biological pathways, including protein phosphorylation, trafficking, lipid metabolism, and mitochondrial function. We demonstrated that GSAP physically interacts with Fe65:APP complex to regulate APP trafficking/partitioning. GSAP is enriched in the mitochondria-associated membrane (MAM) and regulates lipid homeostasis through the amyloidogenic processing of APP. GSAP deletion generates a lipid environment unfavorable for AD pathogenesis, leading to improved mitochondrial function and the rescue of cognitive deficits in an AD mouse model. Finally, we identified a novel GSAP single-nucleotide polymorphism that regulates its brain transcript level and is associated with an increased AD risk. Together, our findings indicate that GSAP impairs mitochondrial function through its MAM localization, and lowering GSAP expression reduces pathological effects associated with AD.

## INTRODUCTION

γ-Secretase activating protein (GSAP) plays an important role in regulating γ-secretase activity and specificity. GSAP selectively modulates of γ-secretase activity toward APP cleavage but not Notch (He et al., 2010; Wong et al., 2019). Depletion of GSAP consistently decreases amyloid-β (Aβ) generation in cells (He et al., 2010; Hussain et al., 2013; Wong et al., 2019). Furthermore, genetic knockdown or pharmacological inhibition of GSAP lowers amyloid plaque deposition and tau phosphorylation in AD mouse models (Chu et al., 2014; Chu et al., 2015; He et al., 2010). Recently, it has been reported that GSAP physically interacts with amyloid precursor protein (APP) to regulate amyloid-β generation (Angira et al., 2019). In addition to increased GSAP levels observed in AD mouse models (Chu et al., 2015), GSAP up-regulation has also been reported in neurodegenerative contexts such as Down Syndrome (Chu et al., 2016), which is obligately associated with Aβ plaque pathology due to triplication of human Chromosome 21 harboring APP (Wiseman et al., 2015). Importantly, several studies have also independently demonstrated that GSAP protein level is significantly increased in postmortem brains of severe AD patients (Chu et al., 2015; Perez et al., 2017; Satoh et al., 2012). Single nucleotide polymorphisms (SNPs) at the GSAP locus have also been identified, and have been shown to correlate with AD diagnosis (Floudas et al., 2014; Zhu et al., 2014). One SNP located in the GSAP promoter region comprises an allele associated with high GSAP expression, which correlates with increased AD risk (Zhu et al., 2014). Together, these studies implicate a pathogenic role for GSAP in AD. Aside from its role in activating γ-secretase activity and APP trafficking/partitioning, little is known with respect to other biological pathways involved in GSAP-dependent AD pathogenesis.

In this study, we identified novel GSAP binding proteins by proteomic analysis and demonstrate that GSAP regulates APP phosphorylation and trafficking/partitioning through physical interactions with the APP binding protein Fe65. We also compared transcriptomic profiles in wildtype (WT) and GSAP knockout (KO) mouse hippocampus by single nuclei RNA sequencing (sn-RNAseq). Pathway enrichment analysis of proteomic and sn-RNAseq datasets concordantly identified overlapping biological pathways associated with GSAP, including protein phosphorylation, trafficking, lipid metabolism, and mitochondrial function. We further demonstrated that GSAP is enriched in the mitochondria associated membrane (MAM) and promotes APP carboxyl terminal fragment (APP-CTF) partitioning into lipid-rafts in favor of Aβ production. We demonstrated that GSAP deletion changed cellular lipid profile and restores impaired memory behavior by novel object recognition tests in the J20 AD mouse model. Finally, a novel SNP was identified and shown to specifically regulate GSAP mRNA expression in human brain; the allele associated with high GSAP expression was found to correlate with AD risk. Taken together, our findings uncover new pathogenic pathways mediated by GSAP, and provide evidence that reducing GSAP levels can attenuate pathogenic events associated with AD pathogenesis.

## RESULTS

### The GSAP protein complex regulates protein phosphorylation, trafficking, lipid metabolism and mitochondrial function

To investigate new players in GSAP function, we identified GSAP binding proteins by two approaches. We first performed co-immunoprecipitation (co-IP) from mouse neuroblastoma N2a cells followed by mass spectrometry (MS) analysis. Using N2a cells to transiently express either HA tagged empty vector (HA-EV) or HA-GSAP plasmids (Fig. 1A), proteins were immunoprecipitated using the HA antibody and proteins specifically enriched in HA-GSAP transfected samples were subjected to KEGG and GO (Gene Ontology) pathway analyses (Fig. 1B,C, S1A). GO pathway analysis suggested that GSAP and its binding protein complex regulate transport, lipid metabolism, and mitochondrial function which are essential pathways altered in AD (Fig. 1B). Interestingly, KEGG pathway analysis demonstrated that GSAP and its binding proteins may be involved in multiple neuronal disorders, including AD (Fig. 1C). Within GSAP binding partners, we identified multiple kinases and phosphatases (Fig. 1D, highlighted in red), in addition to proteins directly involved in trafficking (Fig. 1D, highlighted in blue). A significant number of mitochondrial proteins were also observed in the GSAP interactome (Fig. S1A, highlighted in red). We also assessed the biological function of GSAP using Humanbase, a machine learning-based framework (hb.flatironinstitute.org/gene/54103/ Biological process). In good agreement with our results here, GSAP was predicted to play essential roles in protein transport and phosphorylation regulation (Fig. S1B). Next, we performed yeast two hybrid (Y2H) screening of a human brain cDNA library using the 16 kD carboxy domain of human GSAP (GSAP-16K) as the bait. GSAP-16K is the functional domain responsible for γ-secretase activity regulation (He et al., 2010). We identified 80 proteins that can directly bind GSAP through the 16K domain and also regulate phosphorylation, trafficking, lipid metabolism and mitochondrial function (Fig. S1C).

**Figure 1:**
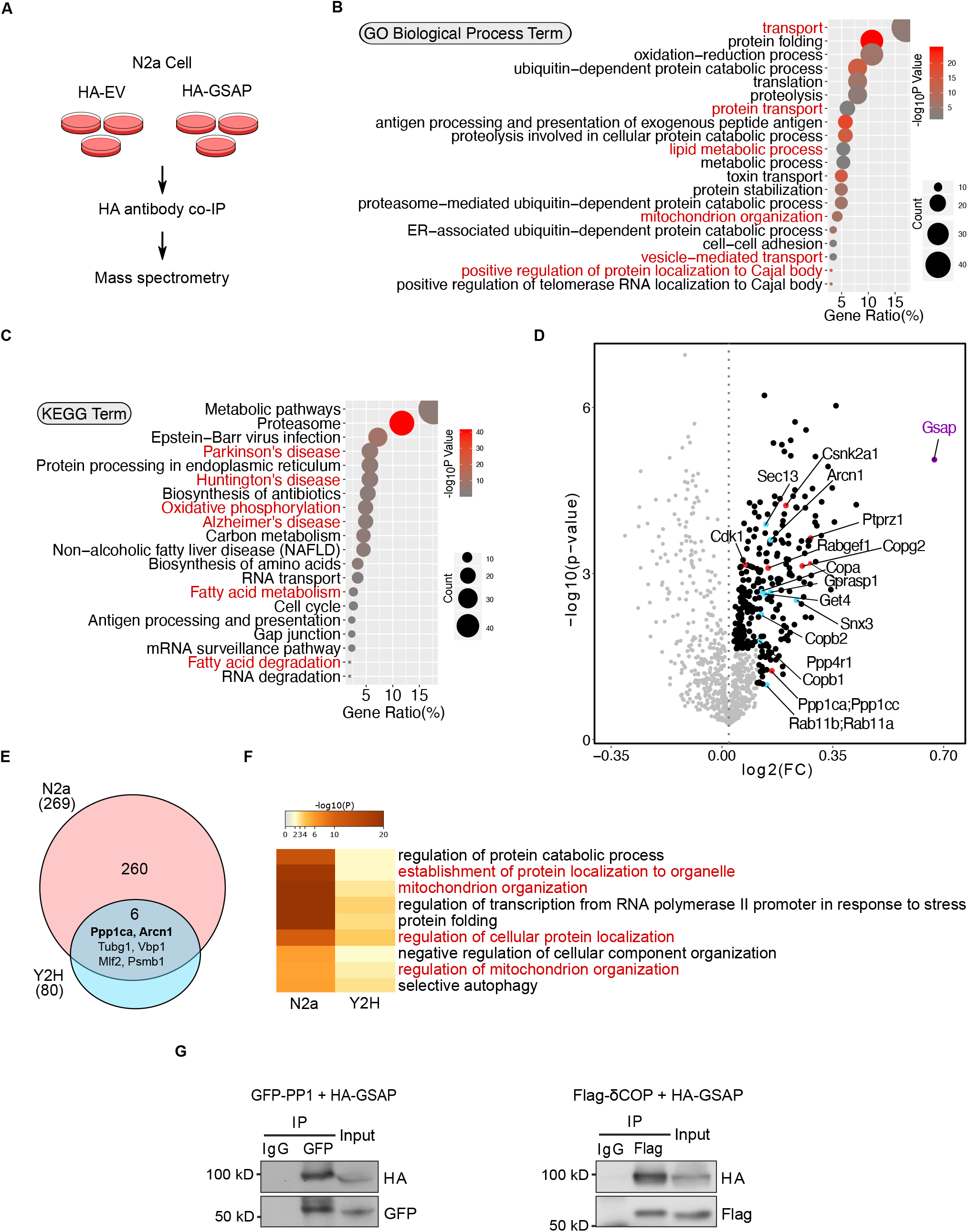
GSAP and its binding proteins are involved in novel biological pathways. **(A)** Schematic of the experimental design to characterize the GSAP interactome. HA-empty vector (EV) was used as a negative control. **(B)** GO pathway enrichment analysis for GSAP binding proteins. Top 20 significantly enriched pathways (p < 0.05) are shown based on p value (dot color) and gene count (dot size). **(C)** KEGG biological process enrichment analysis for GSAP binding proteins. Top 20 significantly enriched pathways (p < 0.05) are shown based on p value (dot color) and gene count (dot size). **(D)** Volcano plot showing differentially enriched proteins (detailed in the methods) in HA-GSAP versus HA-EV co-IP MS experiment in N2a cell. GSAP itself (purple), proteins involved in trafficking (blue), and phosphorylation (red) are highlighted. **(E)** Venn diagram showing overlapped protein between different lists. The circle area is not proportional to the sample size. **(F)** Meta-enrichment analysis of common GO biological pathways shared by three GSAP binding protein lists. **(G)** Co-IP validation of GSAP interaction with PP1 and δ-COP (Arcn1) in HEK293T or N2a cells respectively via transient transfection.

Direct comparison of GSAP binding proteins identified by these two approaches uncovered 6 common proteins (Fig. 1E). Meta-enrichment analysis of shared biological pathways demonstrated that protein trafficking and mitochondria related biological pathways were the top GO pathways shared by these two lists (Fig. 1F). We then validated some of the GSAP binding proteins from the lists, including PP1 (phosphorylation), PHB (mitochondrial function) and δCOP (encoded by the Arcn1 gene; trafficking). We confirmed that GSAP interacts with PP1 (PP1γ encoded by the Ppp1cc gene), PHB and δCOP by co-IP analysis (Fig. 1G, S1D). Notably, we have shown recently that δCOP regulates Aβ production via regulating APP retrograde trafficking, and mutation of δCOP significantly decreases amyloid plaque formation while enhancing cognitive function in an AD mouse model *in vivo* (Bettayeb et al., 2016a; Bettayeb et al., 2016b).

Taken together, our data suggest that the GSAP and its binding proteins play critical roles in regulating protein phosphorylation, trafficking, lipid metabolism and mitochondrial function.

### GSAP directly interacts with the Fe65:APP protein complex and regulates APP phosphorylation and trafficking/partitioning

Previous studies demonstrate that phosphorylation of APP at Thr668 influences APP processing, via a mechanism involving its association with the lipid-raft microdomain (Matsushima et al., 2012). Morevoer, our proteomic analyses revealed an enrichment of components related to protein phosphorylation in the GSAP interactome, suggesting that GSAP may modulate APP Thr668 phosphorylation. We found that GSAP siRNA knockdown significantly increased phospho-Thr668 (pT668) APP levels in N2a cells stably expressing human APP695 isoform (N2a695), with no effect on total APP levels compared to control siRNA transfection (Fig. 2A). Thr668 of APP can be phosphorylated by several kinases to regulate a variety of APP functions (Aplin et al., 1996; Iijima et al., 2000; Standen et al., 2001; Suzuki et al., 1994). In contrast, PP1 is the only protein phosphatase identified to dephosphorylate this site through the recruitment of Fe65, the well-characterized APP binding protein (Rebelo et al., 2013). The Y2H study showed that both PP1 and Fe65 (encoded by the APBB1 gene) directly interact with GSAP (Fig. S1C). In addition to PP1, we also confirmed the interaction of Fe65 and GSAP by co-IP assay in the cells (Fig. 2B), and demonstrated that the GSAP-16K domain was sufficient to bind full length Fe65 (Fig. 2C). We further performed endogenous co-IP experiment using an Fe65 antibody in the human lung carcinoma cell line A549, which has high endogenous expression of both GSAP and Fe65 and showed the co-precipitation of endogenous Fe65 with PP1 and GSAP (Fig. 2D, S1E). Since we observed that knockdown of GSAP decreased APP-CTF association with lipid-rafts (Chang et al., 2020), these results suggest that GSAP regulates APP phosphorylation and partioning through Fe65 interaction.

**Figure 2:**
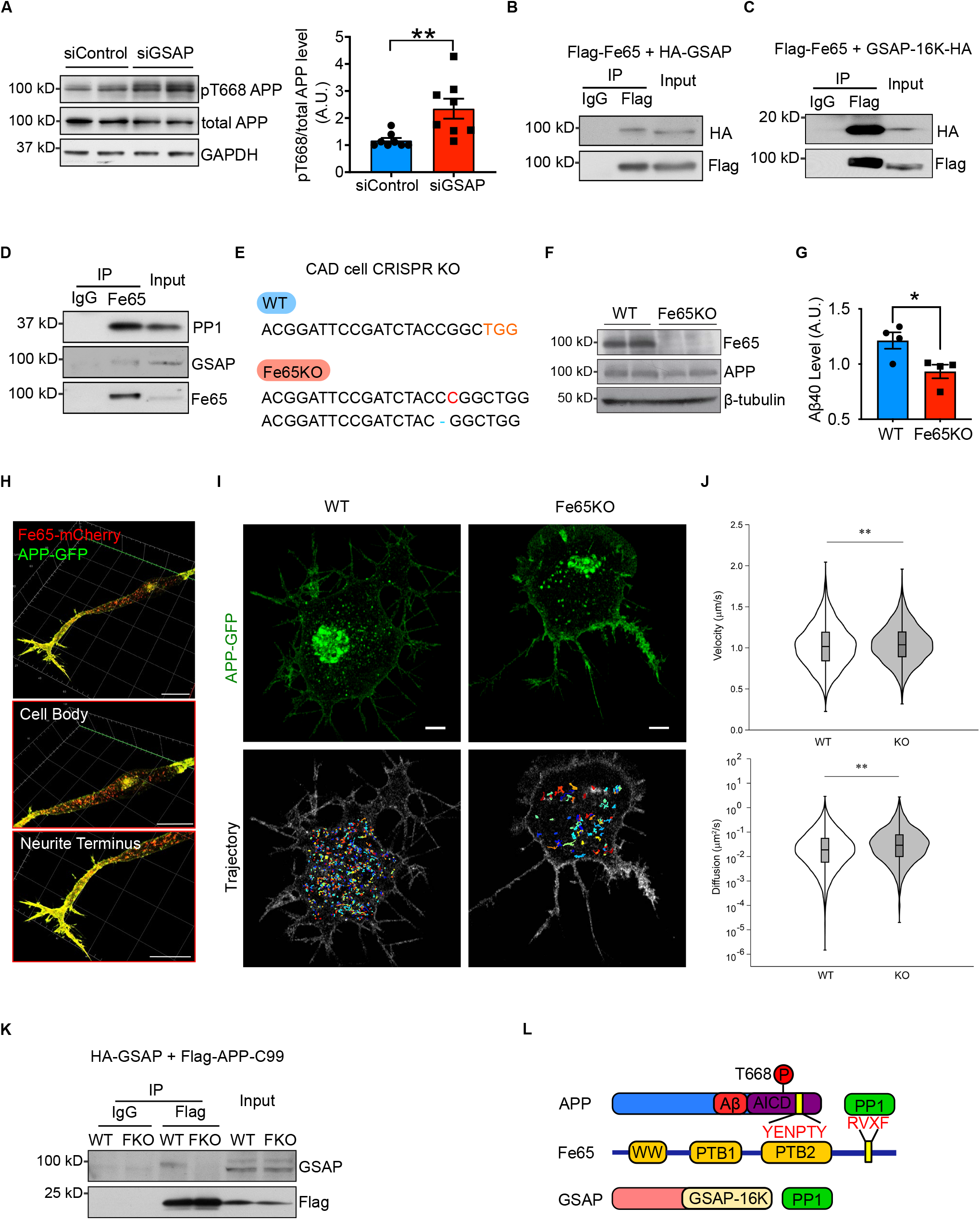
GSAP interacts with Fe65 to regulate APP phosphorylation and trafficking. **(A)** Immunoblot analysis of protein levels in N2a695 cells transfected with control or GSAP siRNA (left panel). Quantification of APP phosphorylation at T668 normalized to total APP level (right panel). Data represent mean ± s.e.m., unpaired t-test, **p < 0.01. **(B)** Co-IP analysis of full length GSAP (HA tagged) interaction with full length Fe65 (Flag tagged) using Flag antibody in HEK293T cells. **(C)** Co-IP analysis of GSAP carboxyl-terminal 16K domain (HA-tagged) co-precipitation with full length Fe65 (Flag-tagged) using a Flag antibody in HEK293T cells. **(D)** Co-IP analysis of endogenous Fe65 interaction with GSAP and PP1 using Fe65 antibody in HEK293T cells. GSAP was detected using an antibody from R&D systems. **(E)** Genomic DNA from CAD WT and Fe65KO cells was isolated and PCR-amplified fragments flanking the CRISPR-Cas9 cleavage site were generated. PCR fragments were cloned into TOPO vector for Sanger sequencing. One base pair insertion (red) and deletion (blue) was identified in Fe65KO CAD cells. **(F)** Immunoblot analysis of proteins from wild-type (WT) and Fe65 knockout (KO) CAD cells. **(G)** Quantification by ELISA of secreted Aβ40 levels produced by WT and Fe65KO cells. Data represent mean ± s.e.m., unpaired t-test, *p < 0.05. **(H)** Representative confocal images of Fe65 (red) and APP (green) localized in differentiated CAD cells. Scale bar: 20 μm. **(I)** Maximum intensity projection of Airyscan Z-stack of WT (top left) and Fe65KO (top right) CAD cells from 95 slices and 0.173 μm step size and generated in Imaris. Scale bar: 5 μm. The images are representative of four independent experiments. WT (bottom left) and Fe65KO (bottom right) trajectories corresponding to the representative time-lapse image series shown in top panel and were reconstructed in MATLAB. Trajectory minimum cut-off time: 10 sec. **(J)** Violin plots showing the velocity (left) and diffusion coefficient (right) distributions of single APP-GFP vesicles in WT and Fe65KO CAD cells. The median value is shown as the horizontal line in the box. The box presents interquartile range. The distributions were compared using the Mann−Whitney U test. (**p < 0.001, WT V_median_ = 1.016 μm/s, KO V_median_ = 1.038 μm/s; WT D_median_ = 0.0187 μm2/s, KO D_median_ = 0.0290 μm2/s). **(K)** Co-IP analysis of GSAP (HA-tagged) with APP-C99 (Flag-tagged) in WT and Fe65KO (FKO) CAD cells. **(L)** Schematic of protein domain interactions within the APP:Fe65:GSAP protein complex.

To further investigate potential biological effects of GSAP:Fe65 interaction, we generated Fe65 knockout (Fe65KO) neuronal tumor CAD cells by CRISPR-Cas9 editing (Qi et al., 1997). Different genomic frame shifts were confirmed on both alleles of the Fe65 gene, which resulted in reduced Fe65 protein levels in Fe65KO cells (Fig. 2E,F). We over-expressed APP and compared Aβ40 levels in WT and Fe65KO CAD cells; consistent with previous observations, Aβ40 levels were reduced in Fe65KO cells (Fig. 2F,G) (Xie et al., 2007). Since GSAP physically interacts with Fe65 and regulates APP intracellular trafficking (Chang et al., 2020), we hypothesized that Fe65 regulates APP intracellular trafficking in a manner similar to GSAP. We first characterized Fe65 and APP sub-cellular localization in CAD cells. Fe65 staining in differentiated CAD cells revealed Golgi-like localization in the cell body and vesicle-like localization at neurite terminals (Fig. 2H). In agreement with previous studies, Fe65 staining showed good overlap with APP (Sabo et al., 2003). We next determined whether Fe65 deletion could affect intracellular APP trafficking by tracking dynamics of single APP vesicles in WT and Fe65KO CAD cells. APP-GFP vesicles were tracked for 1 min under an Airyscan super-resolution microscope at 10 frames/second, and trajectories of each single APP vesicle were analyzed (Fig. 2I). In agreement with our hypothesis, Fe65 regulated APP trafficking dynamics in a fashion similar to GSAP: Fe65KO increased APP vesicle trafficking velocity and diffusivity (Fig. 2J). Since strong binding affinities between Fe65 and APP have been previously established (Radzimanowski et al., 2008), we hypothesized that Fe65 may be required to stabilize GSAP:APP interaction. To test this hypothesis, we compared GSAP and APP interactions in WT and Fe65KO(FKO) CAD cells. HA-tagged GSAP and Flag-tagged APP-C99 (APP C-terminus) proteins were co-expressed in CAD cells and subjected to immunoprecipitation using a Flag antibody. Although, GSAP consistently co-precipitated with APP-C99 in WT cells, GSAP:APP complex formation was dramatically reduced in FKO CAD cells (Fig. 2K). Taken together, this indicates that Fe65 is essential for GSAP:APP interaction, and GSAP-dependent regulation of APP trafficking dynamics.

Fe65 has three well defined protein domains (Bressler et al., 1996; Fiore et al., 1995; Guenette et al., 1996), including the PTB2 domain which directly interacts with the APP AICD domain in the co-crystal structure (Radzimanowski et al., 2008). The RVXF binding motif at the carboxyl terminus of Fe65 may directly interact with PP1 and recruit it to dephosphorylate APP (Rebelo et al., 2013). Together, our data suggest that GSAP is recruited by Fe65 to form a ternary APP/Fe65/PP1 protein complex (Fig. 2L), and demonstrate that GSAP binds Fe65 directly through the GSAP-16K domain and regulates APP phosphorylation.

### GSAP regulates protein phosphorylation, trafficking, lipid metabolism and mitochondrial function in mouse hippocampus *in vivo*

To further investigate the pathogenic function of GSAP in the disease progression, we determined effects of GSAP gene deletion in an AD mouse model. We targeted exons 9 to 11 of the murine GSAP gene locus by flanking loxP sites, and constitutive GSAP knockout (GKO) mice were obtained by crossing GSAP conditional knockout mice with a murine CMV-Cre driver line (Fig. S2A). Genomic PCR and quantitative reverse transcription PCR (qRT-PCR) confirmed successful excision of GSAP exons 9-11, and reduced GSAP mRNA expression (Fig. S2B,C). We next crossed GKO mice with the J20 AD mouse model to investigate effects of GSAP deletion on AD-associated molecular and behavior changes *in vivo* (Mucke et al., 2000).

GSAP expression is broadly detected in various cell types in the brain (Darmanis et al., 2015; Zhang et al., 2014; Zhang et al., 2016). To elucidate the molecular function of GSAP across various cell types in the brain, we performed single nuclei RNA sequencing (sn-RNAseq) on hippocampal tissues obtained from 6-7 month old WT, GKO, J20;WT, and J20;GKO mice (Fig. 3A). A total of 31923 nuclei were clustered based on their transcriptomes and visualized in uniform manifold approximation and projection (UMAP) space. Based on a previous study (Tasic et al., 2016), nuclei were annotated into 7 distinct cell types in an unsupervised manner (Fig. 3B, S3A). The clustering results were validated by visualizing the expression level of known cell type specific marker genes on the violin plot (Fig. 3C). We also visualized the 2-dimensional distribution of nuclei expressing these marker genes in the UMAP space (Fig. S3C). Our data demonstrated that these marker genes were specifically enriched in the annoated cell type clusters, confirming the accuracy of the annotation strategy.

**Figure 3:**
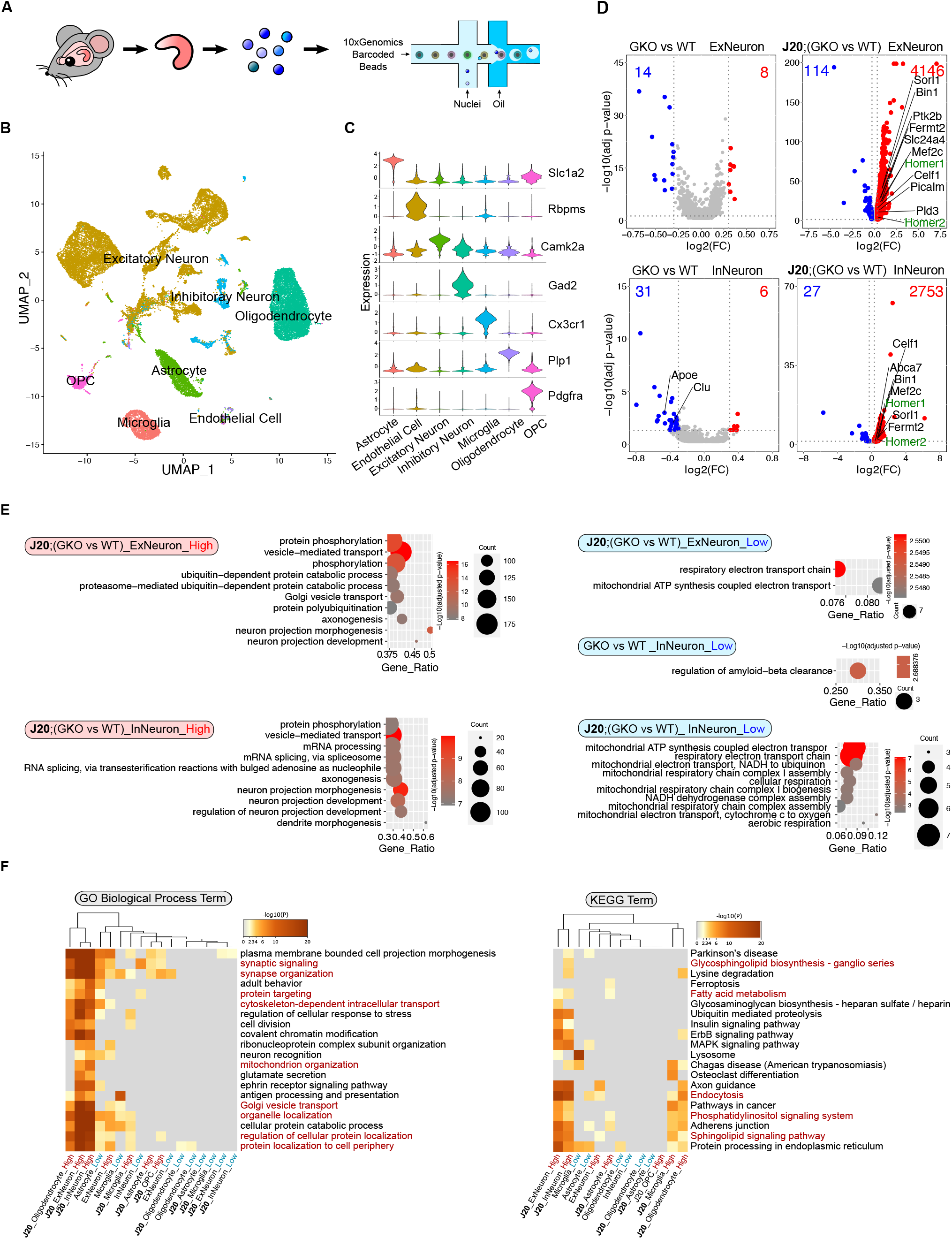
Single nuclei RNA sequencing analysis of GSAP knockout mouse hippocampus. **(A)** Schematic diagram of the experimental design for single nuclei RNA sequencing of mouse hippocampus (WT, GSAPKO, J20;WT and J20;GSAPKO) using the 10X genomics platform. Sequencing data from different genotypes were merged for downstream analysis. **(B)** UMAP plot showing 7 major cell types clustered based on gene expression profile in an unsupervised manner. **(C)** Violin plot showing expression level of representative marker genes from different cell clusters: Slc1a2 (astrocyte; 3220 nuclei), Rbpms (endothelial cell; 213 nuclei), Camk2a (excitatory neuron; 15845 nuclei), Gad2 (inhibitory neuron; 1961 nuclei), Cx3cr1 (microglia; 2210 nuclei), Plp1 (oligodendrocyte; 7187 nuclei), and Pdgfra (oligodendrocyte progenitor cell; OPC; 1287 nuclei). **(D)** Volcano plots showing differentially expressed genes in neuronal clusters comparing WT versus GSAP knockout (GKO) or J20;WT versus J20:GKO. Only genes with significantly expression level change are shown (adjusted p-value <0.05; log2(fold change) < -0.3 or > 0.3). Genes with higher expression level in GKOs are highlighted in red; genes with lower expression level in GKOs are highlighted in blue. AD risk genes are in labeled in black, whereas synaptic genes are labeled in green. **(E)** Gene ontology (GO) biological process enrichment analysis for differentially expressed genes in neuronal clusters. Top significantly changed pathways (up to 10) are shown (adjust p-value <0.05). (**F)** Meta-enrichment analysis of common GO (left panel) and KEGG (right panel) pathways shared by both up-regulated and down-regulated DEGs from all the cell types.

We next sought to determine possible molecular functions of GSAP by examining differentially expressed genes (DEGs) in different cell types in the various mouse genotypes. We compared DEGs in GKO versus WT and J20;GKO versus J20;WT samples. We identified a large number of significant DEGs across cell types in GSAP KO brain, with the exception of endothelial cells which showed comparatively little change. The effect of GSAP deletion on DEGs was largely exacerbated in the J20 mouse model, suggesting that effects of GSAP deletion may be amplified with AD pathogenesis (Fig. 3D, S3B). We first compared GKO DEGs with an AD risk gene list identified from genome wide association studies (GWAS) (Karch and Goate, 2015). Multiple DEGs overlapped with the AD risk gene list in different cell types (Fig. 3D, S3B). In both excitatory and inhibitory neurons, GSAP deletion may confer neuroprotective effects under proteotoxic AD stress since GSAP deletion up-regulates multiple genes previously shown to reduce amyloid-β generation (Pld3, Sorl1, Bin1, and Fermt2). Our data also further supports that loss of function in GSAP KO-induced AD risk genes identified here may contribute to AD pathogenesis (Andersen et al., 2005; Caglayan et al., 2014; Chapuis et al., 2017; Chu et al., 2014; Cruchaga et al., 2014; He et al., 2010; Miyagawa et al., 2016; Ubelmann et al., 2017). Additionally, genes essential for synaptic function (Homer1, Homer2 and Bin1), were significantly up-regulated in both excitatory and inhibitory neurons of J20;GKO mice, suggesting that GSAP depletion may protect synaptic impairment in AD (De Rossi et al., 2020; Shiraishi-Yamaguchi and Furuichi, 2007).

We then characterized biological pathways affected by GSAP deletion. Using GO biological pathway enrichment analysis, we observed that phosphorylation and mitochondrial function were broadly altered in excitatory neurons, inhibitory neurons, oligodendrocytes, microglia and astrocytes (Fig. 3E and S3D). Moreover, we observed enrichment of pathways related to vesicle-mediated transport in neurons and oligodendrocytes with GSAP deletion (Fig. 3E and S3D). We also performed meta-enrichment analysis using DEG lists from all the cell types to identify common biological pathways affected by GSAP depletion. Trafficking (GO term), mitochondrial function (GO term), and lipid metabolism (KEGG term) were among the top shared biological pathways in a variety of cells affected by GSAP depletion (Fig. 3F). Notably, these biological pathways highly overlap with the GSAP functional pathways identified via proteomics, confirming the robustness of our analyses (Fig. 1B,C).

### Characterization of GSAP function in excitatory neurons

Neurons are the major source for Aβ production (Zhao et al., 1996). Having observed that GSAP regulates Aβ production and has the highest expression level in human neurons (Darmanis et al., 2015; Zhang et al., 2016), we then focused on characterizing GSAP function in excitatory neurons, which represents the largest cell population in hippocampus. Since co-expressed genes often work in the same function cluster, we first applied weighted gene co-expression analysis (WGCNA) on DEGs from excitatory neurons to identify gene modules that function as groups (Zhang and Horvath, 2005). We determined the correlation between WGCNA gene modules and mouse genotypes and identified the light cyan WGCNA module had the strongest correlation with genotype (Fig. S3E). GO pathway analysis demonstrated the light cyan module represented the mitochondrial function category (Fig. 4A). Our data suggest that GSAP knockout and/or amyloidogenesis mainly affect mitochondrial function in excitatory neurons.

**Figure 4:**
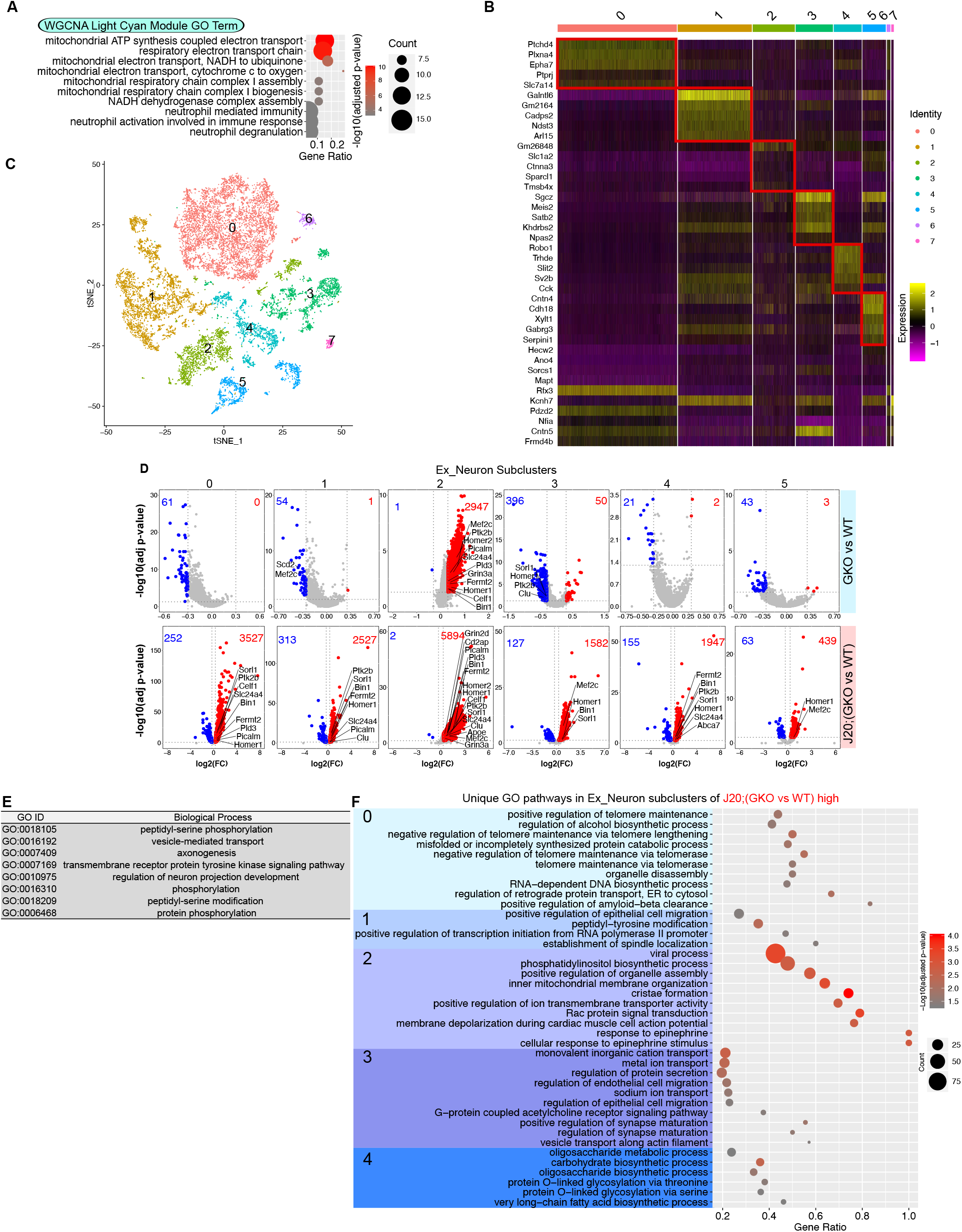
Characterization of GSAP function in excitatory neurons. **(A)** GO biological process enrichment analysis for genes enriched in the WGCNA lightcyan module of excitatory neurons. Top significantly changed pathways (up to 10) are shown (adjust p-value <0.05). **(B)** Heatmap showing the expression level of the top 5 differentially enriched genes in each excitatory neuron sub-cluster. **(C)** t-SNE plot depicting the excitatory neuron cluster, which is divided into 8 sub-clusters in an unsupervised manner. **(D)** Volcano plots showing differentially expressed genes in excitatory neuron sub-clusters comparing GKO versus WT or J20;GKO versus J20:WT. Genes with significant changes in expression levels are shown (adjusted p-value <0.05; log2(fold change) < -0.3 or > 0.3). Up-regulated genes in GKO are highlighted in red; genes down-regulated in GKO are highlighted in blue. **(E**,**F)** GO biological process pathways analyses were performed with up-regulated genes comparing J20;GKO versus J20;WT mice. GO pathways shared by all 5 neuron sub-clusters are shown in (E); GO pathways uniquely over-represented in specific clusters are shown in (F).

Since substantial heterogeneity in gene expression was observed within excitatory neurons, we divided excitatory neurons into different sub-clusters. We classified 8 sub-clusters in excitatory neurons based on nuclear transcriptomic profile (Fig. 4B,C). Significant DEGs were identified from 5 of these sub-clusters (Fig. 4D). Sub-cluster 2 from excitatory neurons exhibited the greatest effect of GSAP depletion, exemplified by the highest number of up-regulated genes even in GKO mice. These results suggest that GSAP mainly functions in the sub-cluster 2 excitatory neurons under physiological conditions, and this effect is potentially exacerbated in the entire excitatory neuron cluster under pathogenic conditions. Comparing J20;GKO with J20;WT, analysis of DEGs suggest that GSAP functions similarly in all 5 sub-clusters in terms of amyloidogenesis and synaptic functional genes regulation (Fig. 4D). To elucidate functional pathway changes in the 5 sub-clusters, we searched for GO biological pathways enriched in up-regulated DEGs in J20;GKO compared to J20;WT mice and identified shared and unique functional pathways. 8 GO terms showed consistent over-representation in all 5 sub-clusters (Fig. 4E). These results demonstrate that GSAP has a general function in the regulation of phosphorylation and trafficking across excitatory neuron sub-types. Unique pathways in individual sub-clusters suggest that GSAP may specifically regulate telomere lengthening in cluster 0 neurons, cell migration in cluster 1 neurons, lipid homeostasis and mitochondrial function in cluster 2 neurons, ion transport and synapse maturation in cluster 3 neurons, and oligosaccharide metabolism in cluster 4 neurons (Fig. 4F).

### GSAP regulates lipid metabolism and mitochondrial function in the MAM

We next sought to investigate the underlying mechanism by which how GSAP regulates mitochondrial function in neuronal cells. Numerous studies have shown that the mitochondria associated membrane (MAM) is an essential hub for the regulation of lipid homeostasis, mitochondrial function, and AD pathogenesis (Area-Gomez et al., 2018; Area-Gomez et al., 2012). Specifically, APP is partitioned and processed in the MAM to generate its C99 fragment and Aβ production, which in turn have detrimental effects on lipid homeostasis and mitochondrial function (Del Prete et al., 2017; Pera et al., 2017). Multiple lines of evidence from our data suggest that GSAP may be enriched in MAM and regulate lipid homeostasis and mitochondrial function through MAM. Our proteomic and genomic data concordantly suggest that GSAP regulates trafficking, lipid metabolism, and mitochondrial function. Secondly, our preliminary data suggest that GSAP protein can interact with phospholipids and mitochondrial enriched cardiolipin (Fig. S4A,B). GSAP interacts with the mitochondrial protein Phb, which was observed in MAM and proposed to regulate lipid homeostasis (Fig. S1D) (Osman et al., 2009; Zhang et al., 2011). Furthermore, enrichment in the MAM fraction has been reported for several GSAP binding proteins including APP, Psen1, Fe65, Arcn1, Copa, Copb2, Htra2, Acsl1, Hspa9, and ER Lipid Raft Associated 2 (Erlin2) (Fig. S1A) (Ma et al., 2017; Schon and Area-Gomez, 2013; Volgyi et al., 2018). Similar to GSAP, the MAM protein Erlin2 binds Psen1 and regulates γ-secretase activity towards APP processing with little or no effect on Notch (Browman et al., 2006; Teranishi et al., 2012).

Since we observed Erlin2 interaction with GSAP in our proteomics data, we first validated their interaction using co-IP. Flag tagged Erlin2 pulled down HA tagged GSAP and endogenous Psen1 (Fig. 5A). We next directly tested whether GSAP is enriched in MAM by cell fractionation (Lewis et al., 2016). To distinguish subcellular ractions, we first analyzed proteins previously shown to be located in MAM. Consistent with previous studies, we observed both Phb and Calnexin in the MAM fraction (Horner et al., 2015; Zhang et al., 2011). In the same assay, we found that GSAP was enriched in MAM (Fig. 5B). Next, we analyzed ER-mitochondria association directly using electron microscopy (EM). We observed close ER-mitochondria (ER-Mito) contacts in both WT and previously established GKO SHSY-5Y cells (Wong et al., 2019) (Fig. 5C). Quantification of ER-Mito contact demonstrated that the proportion of mitochondria with ER contact significantly increased in GKO cells, whereas ER-Mito contact length significantly decreased in GKO cells (Fig. 5D,E). Specifically, a majority of the ER-Mito contacts were very short in GKO cells (< 100 nm), suggesting GSAP is an essential regulator for ER-Mito interaction (Fig. 5F).

**Figure 5:**
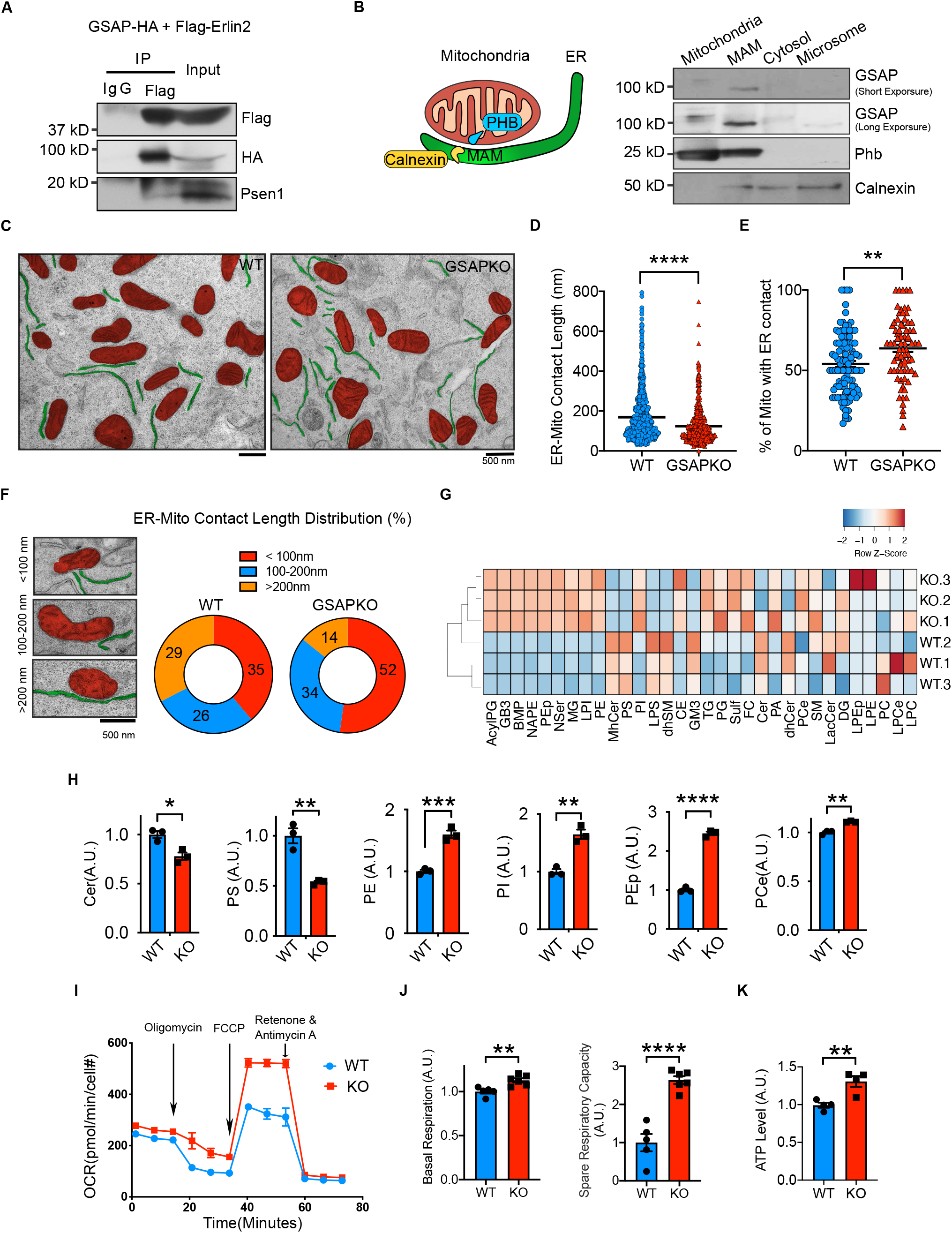
GSAP regulates lipid metabolism and mitochondrial function through the MAM. **(A)** Co-IP analysis of GSAP (HA-tagged) interaction with Erlin2 (Flag-tagged) and Psen1 using an HA antibody in N2a cells. **(B)** Diagram of mitochondria associated ER membrane (MAM), left panel. Equal amount of protein from different fractions of N2a695 cells were loaded into each lane for SDS-PAGE and western blot analysis, right panel. **(C)** Representative electron microscopy (EM) images of WT and GSAP KO SHSY-5Y cells. Mitochondria (red) and ER (green) are highlighted. Scale bar: 500 μm. **(D)** ER-mitochondria (ER-mito) contact length are quantified on the ER side in WT (n=106) and GSAP KO cells (n=76). Data represent mean ± s.e.m., unpaired t-test, ****p < 0.0001. **(E)** The proportion of mitochondria with ER contact is quantified in WT (n=106) and GSAP KO cells (n=76). Images were analyzed in a blinded manner. Data represent mean ± s.e.m., unpaired t-test, **p < 0.01. **(F)** Representative EM images of 3 categories of ER-mitochondria contact based on contact length (left panel). Proportion of ER-Mito contact length in each category is quantified in WT (n=106) and GSAP KO cells (n=76). **(G)** Heatmap showing levels of different lipid subclasses in WT and GSAP KO SHSY-5Y cells by lipidomic analysis. **(H)** Ceramide (Cer), phosphatidylserine (PS), phosphatidylethanolamine (PE), phosphatidylinositol (PI), plasmalogen phosphatidylethanolamine (PEp) and ether phosphatidylcholine (PCe) levels were quantified based on lipidomics analysis. Data represent mean ± s.e.m., unpaired t-test, *p < 0.05, **p < 0.01, ***p < 0.001, ****p < 0.0001. **(I)** Oxygen consumption rate (OCR) of WT and GSAP KO cells were measured in real time by Seahorse assay. Data were normalized to cell count and represent mean ± s.e.m., **(J)** Basal OCR and spare respiration capacity (SRC) were compared between WT and GSAP KO cells. Data represent mean ± s.e.m., unpaired t-test, **p < 0.01, ****p < 0.0001. **(K)** Total intracellular ATP was compared between WT and GSAP KO cells cultured in media without glucose. Data represent mean ± s.e.m., unpaired t-test, **p < 0.01.

To evaluate if GSAP regulates lipid homeostasis, we systematically assessed changes of different lipid classes via lipidomic analysis in WT and GKO SHSY-5Y cells. We noticed large changes in different classes of lipid levels after GSAP knockout (Fig. 5G, S4A). Similar to γ-secretase inhibition (Area-Gomez et al., 2012), GSAP knockout increased phosphatidylethanolamine (PE), confirming effects of GSAP deletion on MAM function (Fig. 5H). Notably, we also observed that GSAP knockout decreased cellular ceramide (Cer) and phosphatidylserine (PS) levels, and increased phosphatidylinositol (PI), plasmalogen phosphatidylethanolamine (PEp) and ether phosphatidylcholine (PCe) levels (Fig. 5H, S4A). The cellular lipid profile changes showed opposite direction of AD pathogenesis (Kosicek and Hecimovic, 2013), suggesting that GSAP knockout may reverse the cellular lipid environment which facilitates AD pathogenesis.

Amyloidogenic processing of APP in MAM, the intracellular lipid-rafts like domain, has been demonstrated to directly regulate lipid homeostasis and mitochondrial function. Specifically, MAM accumulation of APP-C99 triggers the up-regulation of Cer level, which leads to the mitochondrial oxidative phosphorylation defects in AD (Area-Gomez et al., 2018; Area-Gomez et al., 2019). Interestingly, we have observed that GSAP is enriched in lipid-raft microdomains; and knockdown of GSAP decreases APP-CTF association with lipid-rafts (Chang et al., 2020). Importantly, Cer level is significantly decreased after GSAP knockout (Fig. 5H). We therefore tested if GSAP knockout affected mitochondria bioenergetic capacity. We measured mitochondrial oxygen consumption rate (OCR) in WT and GSAP knockout cells using the Seahorse assay (Fig. 5I). Compared to WT, GSAP knockout significantly increased both basal respiration and spare respiratory capacity (SRC) (Fig. 5I,J), which is critical for neuronal survival under cellular stress (Desler et al., 2012). Consistently, total ATP levels were increased in GSAP knockout cells compared to WT (Fig. 5K). Our results suggest that GSAP deficiency improves mitochondrial bioenergetics, which showed deficits early in AD pathogenesis (Terada et al., 2020; Yao et al., 2009). In summary, our data demonstrate that GSAP is enriched in the MAM and regulates lipid homeostasis and mitochondrial function in the MAM.

### Knockout of GSAP rescues novel object recognition behavior in the J20 AD mouse model

To further investigate the biological function of GSAP, we assessed whether GSAP impacts cognitive functions in an AD mouse model by examining alterations in mouse behavior in J20;GKO mice. Since J20 mice exhibit major cognitive deficits at 5-7 months of age, we used 6-month-old J20;WT and J20;GSAP-/-(J20;KO) mice for behavioral analysis (Harris et al., 2010). We did not observe weight differences at 6 months of age (Fig. S2D). Novel object recognition tests were used to evaluate whether GSAP affects recognition memory in AD mice (Fig. 6A). During the re-habituation phase, no difference in travel distance was observed, indicating comparable locomotor activity with GSAP deletion (Fig. 6B, left panel). During the choice phase, J20;WT mice spent a similar amount of time exploring novel and old objects, whereas J20;KO mice spent significantly more time with the novel object (Fig. 6B, middle panel). Preference index showed that J20;GKO mice had significantly greater preference towards the novel object (Fig. 6B, right panel). These results indicate that GSAP knockout restores the recognition memory deficits in the J20 AD mouse model (Mucke et al., 2000).

**Figure 6:**
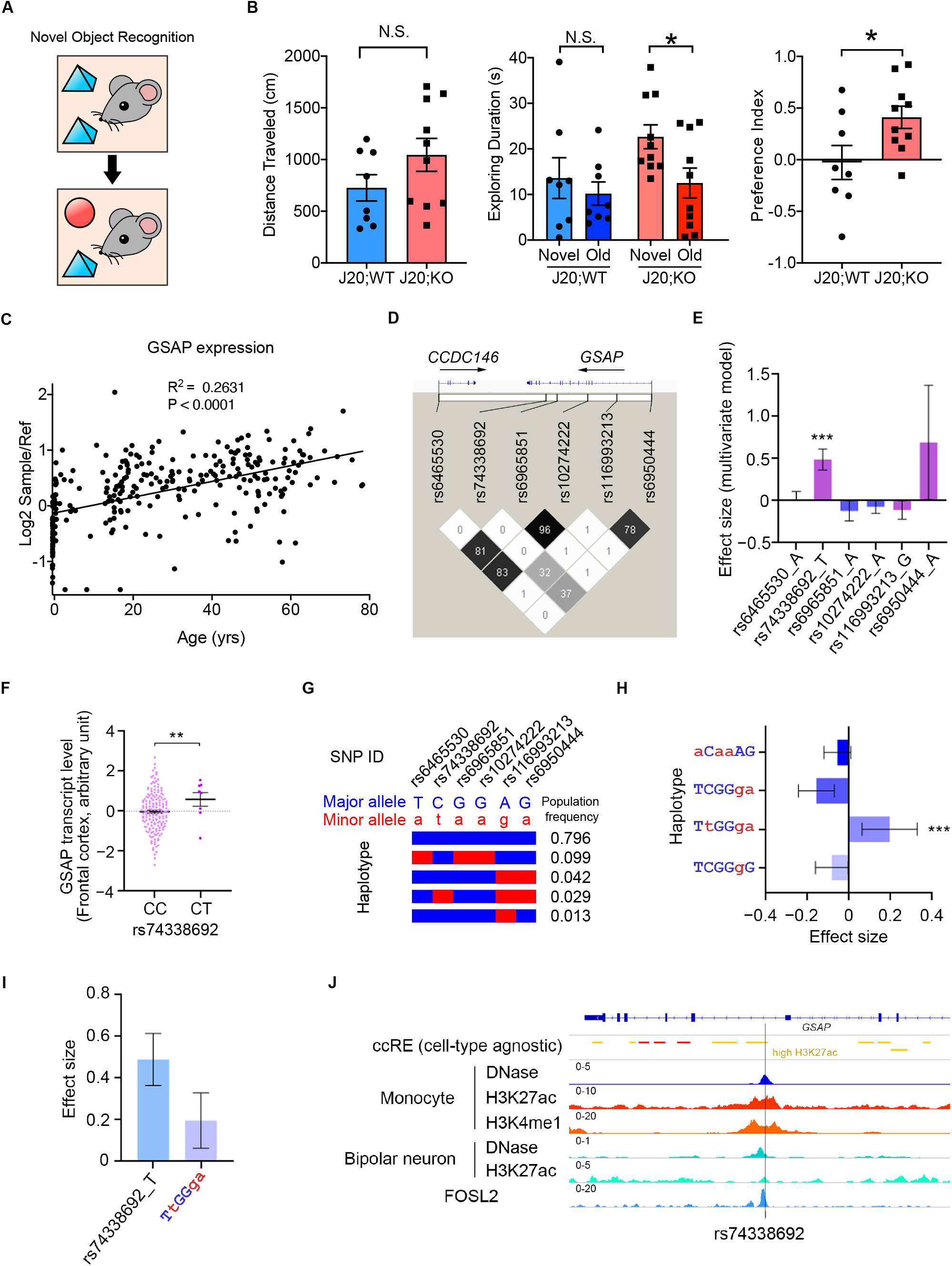
GSAP is involved in the pathogenesis of Alzheimer’s disease in the J20 mouse model and humans. **(A)** Diagram representing the novel object recognition test. **(B)** Memory behavior in 6-month-old J20;WT (5M,3F) and J20;GSAP knockout (J20;KO,4M,6F) were evaluated by novel object recognition test. Total distance traveled during the re-habituation phase was quantified (left panel). Exploration time for old and novel objects during the choice phase were quantified in both genotypes (middle panel). Preference index = (Time novel - Time familiar)/(Time novel + Time familiar)(right panel). Data represent mean ± s.e.m., unpaired t-test, *p < 0.05, N.S. not significant.). **(C)** Human GSAP transcript levels are up-regulated with age. Data were obtained from BrainCloud.**(D)** linkage disequilibrium analysis of GSAP AD risk variants which located in the cis-regulatory elements. The color map corresponds to the pairwise R^2^ values between variants, with values of R^2^ also marked in the plot. **(E)** Association between GSAP AD risk variants and brain GSAP transcript level. The plot shows the effect sizes and corresponding standard errors obtained from the meta-analysis of the results from different brain regions. Data represent effect size ± s.e. ***p < 0.001. **(F)** Association between rs74338692 and GSAP transcript level in the frontal cortex. (n = 167 and 8, for CC and CT, respectively). Data represent mean ± s.e.m. **p < 0.01. **(G)** Haplotype structure defined by the GSAP AD risk variants. Each bar represents a haplotype defined by the minor (red) or major (blue) alleles of six selected variants, with the population frequency (European population from 1000 Genomes Phase3 data; n = 503) marked on the right side of corresponding haplotypes. **(H)** Association between GSAP haplotypes and brain GSAP transcript level. The uppercase (blue) and lowercase (red) letters denote the major and minor alleles of the corresponding variants. The plot shows the effect sizes and corresponding standard errors obtained from the meta-analysis of the results from different brain regions. Data represent effect size ± s.e. ***p < 0.001. **(I)** Comparison between rs74338692 and rs74338692-associaited haplotypes for their associations with brain GSAP transcript level. The plot shows the effect sizes and corresponding standard errors obtained from the meta-analysis of the results from different brain regions. Data represent effect size ± s.e. **(J)** Database evidence suggest the potential regulatory roles of rs74338692. From upper to lower panel: ccRE: Cell type-agnostic annotation for cis-regulatory elements from SCREEN database (hg19 version); Yellow color denotes the cis-regulatory regions with high H3K27ac signal. DNase: the normalized signal of DNase I hypersensitive sites sequencing data in different cell types. H3K4me1, the normalized signal of H3K4me1 ChIP-seq data in monocytes. H3K27ac, the normalized signal of H3K27ac ChIP-seq data in different cell types. FOSL2: the normalized signal of FOSL2 ChIP-seq in HepG2 cells. The heights of each track were labelled on the upper-left corner of corresponding tracks, with cell type information labelled on the left of the tracks.

### Transcriptional regulation of GSAP correlates with human aging and AD

Since aging is the highest risk factor for AD, we first analyzed GSAP mRNA expression across different human brain regions at different ages. Data from both BrainCloud and PsychENCODE independently provided evidence that GSAP transcripts increased with age, across varying human brain regions (Fig. 6C, S5A). Increased GSAP mRNA expression with age supports previous findings that GSAP protein levels are significantly elevated in AD patient brain with severe pathology and cognitive deficits (Perez et al., 2017; Satoh et al., 2011; Zhu et al., 2014). These results indicate GSAP may contribute to human AD during aging.

Multiple GSAP SNPs have been previously associated with AD (Floudas et al., 2014; Zhu et al., 2014). A candidate genetic study has identified the potential association between GSAP promoter variants and AD risk in a Chinese AD cohort (Zhu et al., 2014). To conduct a comprehensive analysis of AD genetic risk for GSAP, we investigated the results from the up-to-date AD meta-analysis (Jansen et al., 2019). By querying the summary statistics obtained from the AD meta-analysis generated from ∼ 450k individuals (n = 455,258), several non-coding variants (p < 0.05) resided in the GSAP locus that exerted AD risk were identified (Fig. 6D, S5B). Six of the identified variants were located in the annotated cis-regulatory elements, with the majority of them being found in the regions with transcription factor binding events (Fig. S5C). To investigate their possible effects on GSAP level, association analysis was conducted between the identified variants and the brain GSAP transcript level, with only rs74338692 exerting significant association after meta-analysis summarizing results from 13 brain regions (p < 0.001; Fig. 6E; Fig. 6F as a demonstration for association between rs74338692 and GSAP expression in the frontal cortex).

As rs74338692 is in moderate linkage disequilibrium with other SNPs residing in the regulatory regions (R^2^ > 0.3), it is possible that a haplotype defined by the risk alleles of these variants may exert a greater effect on modulating GSAP expression than rs74338692 alone. Haplotype analysis regarding the six SNPs were conducted, and there was only one haplotype (TtGGga) harboring the AD risk allele rs74338692 (Fig. 6G). Further association analysis again revealed that only this haplotype, TtGGga, was significantly associated with brain GSAP transcript level (p < 0.001; Fig. 6H). This haplotype did not exert a higher effect size for modulating GSAP brain transcript level as compared to the rs74338692 alone (Fig. 6I), suggesting that rs74338692 might be major genetic factor that modulate expression of GSAP in the brain. Subsequent investigation of epigenetic profiles in rs74338692-associated genomic region revealed its overlap with the high H3K27ac signal, a marker for enhancer activity, in a cell type-agnostic manner. By specifically examining the epigenomic profiles of monocytes (with high GSAP expression), additional signals for regulatory regions or enhancer activity, including DNase, H3K4me1, and H3K27ac, were again observed in the rs74338692-associated genomic region. Notably, this region also exerted regulatory property in neuronal cells types, including DNase and H3K27ac. Moreover, transcription factor binding activity was also observed in this region, as suggested by the ChIP-seq results of FOSL2, a subunit of AP-1 transcription factor complex (Fig. 6I). In summary, our results suggest a potential regulatory function of the rs74338692-harbored genomic region, which might be the underlying mechanism of how the AD risk GSAP variant rs74338692 may lead to elevated GSAP level in the brain.

### DISCUSSION

Clinical AD trials have so far tested two γ-secretase inhibitors (GSI; Semagacestat and Avagacestat), the failure of which highlighted the importance of developing selective γ-secretase modulators (GSM) (Coric et al., 2012; Doody et al., 2013; Karran and Hardy, 2014; Mekala et al., 2020; Nie et al., 2020). We previously implicate GSAP as an attractive target for selective γ-secretase modulation based on two related mechanisms: 1) GSAP specifically regulates γ-secretase catalytic activity towards APP through modulating PS1 conformation, and 2) GSAP regulates APP trafficking and partitioning into lipid-raft microdomain, where γ-secretase is enriched (Chang et al., 2020; He et al., 2010; Wong et al., 2019). Using proteomics and sn-RNAseq, our current work unbiasedly uncovered potential molecular functions of GSAP in the regulation of protein phosphorylation, trafficking, lipid metabolism and mitochondrial function. Many of these pathways are directly affected by γ-secretase and dysregulated in late onset AD (LOAD), further supporting that selective γ-secretase modulation via GSAP could be beneficial in AD treatment.

GSAP is broadly expressed in various brain cell types, and shows highest expression in neurons in humans (Darmanis et al., 2015; Zhang et al., 2014; Zhang et al., 2016). GSAP deletion significantly changes transcriptomic profiles in almost all cell types including neurons. Pathway analysis of our proteomic and genomic data concordantly revealed protein phosphorylation and trafficking as the top GSAP regulated biological pathways shared by different cell types. Previous studies have extensively shown that neuronal APP trafficking is regulated by protein phosphorylation, and represents one of the most essential pathways in AD pathogenesis (Haass et al., 2012). Recently, we observed that APP trafficking and partitioning in neuronal cells is regulated by GSAP (Chang et al., 2020), which may occur through the novel GSAP/Fe65/APP/PP1 protein complex described here. Since conflicting results have been reported with respect to the direct interaction of GSAP and APP, our current data favor a molecular model where Fe65 recruits GSAP:PP1 to dephosphorylate APP and regulate its trafficking and partitioning to lipid rafts (Angira et al., 2019; Deatherage et al., 2012; Savolainen et al., 2015). It would be interesting to further explore intermolecular interactions within the GSAP:Fe65:APP:PP1 protein complex and investigate how this complex regulates APP trafficking and partitioning. Additionally, our sn-RNAseq data also indicate that GSAP may regulate microglia activation, which needs further exploration.

Lipid metabolism and mitochondrial function are also the top biological pathways regulated by GSAP. It has been established that mitochondria dysfunction contributes to AD pathogenesis. (Guo et al., 2020; Wang et al., 2020). Notably, mitochondrial function had the strongest correlation with GSAP knockout and/or amyloidogenesis in excitatory neurons. Previous work suggested that the MAM is a central hub for lipid metabolism and mitochondrial function regulation (Area-Gomez et al., 2018). It was demonstrated that the amyloidogenic processing of APP in the MAM is responsible for the dysregulation of lipid metabolism (Del Prete et al., 2017; Pera et al., 2017). Interestingly, we demonstrated that GSAP is localized in the MAM, the intracellular lipid-raft like domain, and knockdown of GSAP decreases APP-CTF accumulation in the lipid-rafts and decreases Aβ production. Hence GSAP may function through modulating both APP partitioning to the MAM and γ-secretase activity in the MAM to regulate lipid metabolism and mitochondrial function. Indeed, GSAP depletion decreases ER-mito contacts, which were shown to be increased in different models of AD pathogenesis (Area-Gomez et al., 2012; Del Prete et al., 2017; Hedskog et al., 2013; Martino Adami et al., 2019). Notably, GSAP depletion significantly decreases ceramide level, a known apoptogenic mediator and important neurodegeneration regulator, which is commonly increased in human AD brain (Jana et al., 2009; Kolesnick and Kronke, 1998; Kosicek and Hecimovic, 2013). Cellular ceramide level can be regulated by various amyloidogenic products of APP: APP-C99 accumulation in the MAM increases ceramide synthesis (Pera et al., 2017), different forms of Aβ also induce ceramide synthesis and cell death in neurons and glial cells (Ayasolla et al., 2004; Lee et al., 2004; Malaplate-Armand et al., 2006; Zeng et al., 2005). In addition to ceramide, GSAP depletion reverses the cellular lipid environment in the opposite direction of AD pathogenesis. Depletion of GSAP increases PE, PI, PEp, and PCe levels, and decreases PS levels. Human AD brain showed consistent decreases in PE, PI, and PEp compared to control (Kosicek and Hecimovic, 2013). Moreover, PCe was decreased in the PS1/APP AD mouse model in the cortex compared to control (Ojo et al., 2019) and PS can mediate synaptic pruning by microglia as an “eat-me” signal (Scott-Hewitt et al., 2020). It was also demonstrated that PEp can reduce γ-secretase activity for Aβ production, preventing neuronal death (Su et al., 2019). We also observed that GSAP knockout also decreases the level of lysophosphatidylserine, the up-regulation of which can promote neurodegeneration through microglia (Blankman et al., 2013).

Lipid metabolism significantly contributes to the mitochondrial function. Previous work has demonstrated that the increase in cellular ceramide may be the major cause of subsequent mitochondrial dysfunction (Pera et al., 2017). Moreover, PE deficiency can also impair mitochondrial bioenergetic function (Tasseva et al., 2013). In agreement with this idea, the lipid profile changes after GSAP depletion may largely contribute to the improvement of mitochondrial bioenergetic function. It is of particular interest that GSAP depletion significantly increases mitochondrial spare respiratory capacity (SRC). SRC is thought to generate extra energy supply to maintain cellular function, especially under stress (Sansbury et al., 2011). Consistently, enhanced SRC promotes cell survival, whereas reduced SRC may contribute to cell death (Nickens et al., 2013; Yadava and Nicholls, 2007). In AD, deficiency of SRC was shown to contribute to neuropsychological changes (Bell et al., 2020). Since mitochondrial bioenergetic function is one of the early deficits in AD (Terada et al., 2020; Yao et al., 2009), reduction of GSAP level may delay the pathogenesis of AD. Lastly, GSAP also interacts with several components of endoplasmic reticulum-associated degredation (ERAD) machinery, a protein quality control mechanism that regulates mitochondrial function through MAM and is critical in AD pathogenesis (Zhou et al., 2020; Zhu et al., 2017). Further studies will be needed to characterize the functional interactions between ERAD and GSAP in AD pathogenesis.

Similar to IFITM3, the newly identified γ-secretase modulatory protein (Hur et al., 2020), GSAP level is significantly induced by inflammatory responses and upregulated by ageing and AD pathogenesis in humans. Its expression is induced by LPS and IFNγ in macrophages (Orecchioni et al., 2019), and by LPS alone in primary microglia cells (He et al., 2018). Our results indicate that GSAP deletion restores cognitive function in mice and human GSAP plays important role in the pathogenesis of AD. Furthermore, the GSAP homologue in drosophila was found to genetically interact with the intermediate early transcription factor AP-1, and consequently regulate neuronal AP-1 function (Franciscovich et al., 2008). Since AP-1 function is critical for neuroplasticity, learning and memory, it would be interesting to investigate the interaction of GSAP with AP-1 in mammalian system and further determine its function in learning and memory (Gallo et al., 2018).

In summary, our work indicate that GSAP regulates lipid metabolism, mitochondrial bioenergetic function in the MAM through modulating both APP partitioning and γ-secretase catalytic activity, suggesting GSAP is a pathogenic component of human AD and exacerbates AD phenotypes in AD mice. Thus, reducing GSAP levels may ameliorate cognitive deficits in AD (Fig. 7).

**Figure 7:**
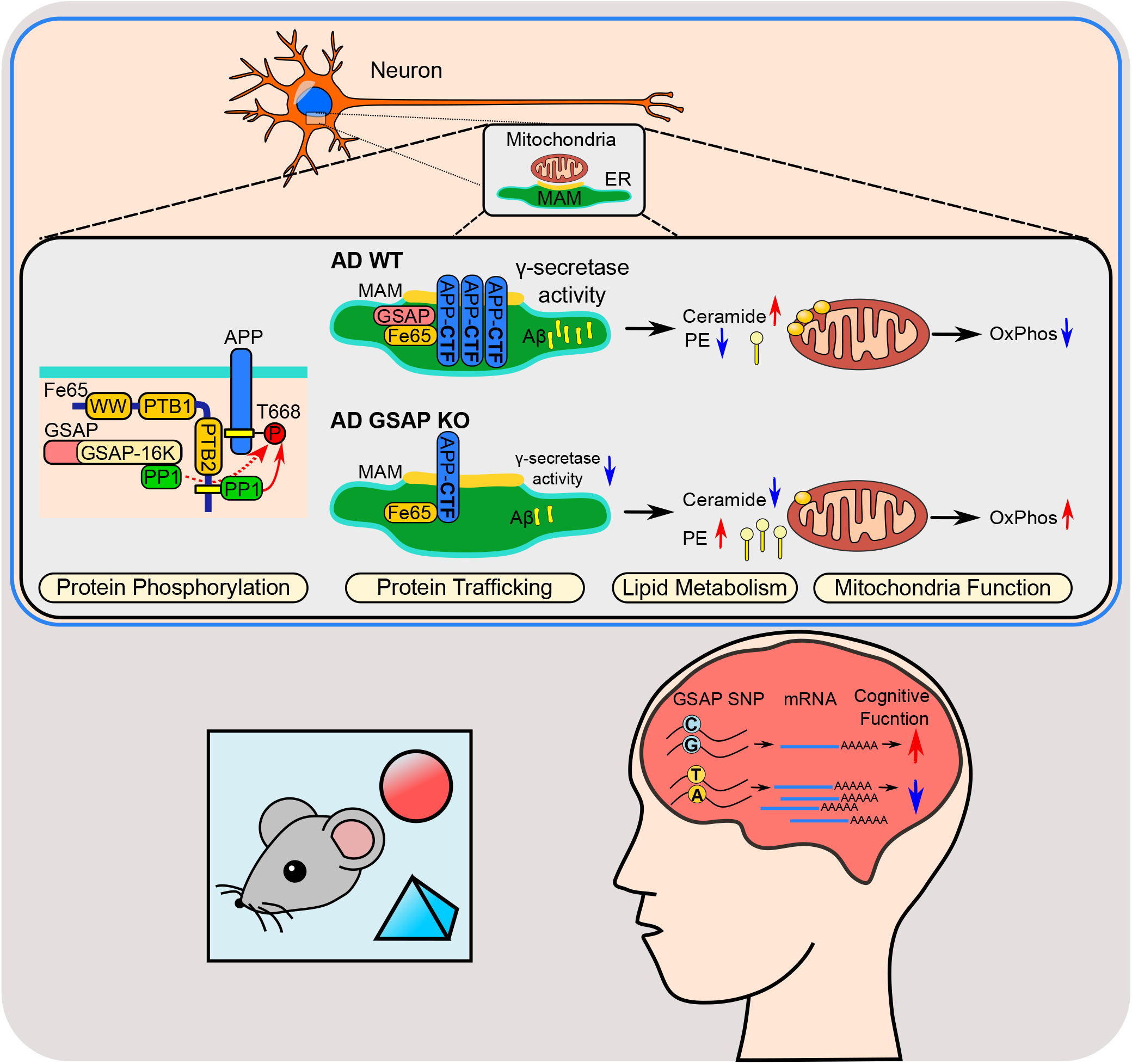
Summary model. GSAP is involved in late onset AD-related pathways including protein phosphorylation, trafficking, lipid metabolism, and mitochondrial function. In neurons, GSAP forms a complex with Fe65:PP1:APP to regulate APP phosphorylation; depletion of GSAP decreases APP-CTF partitioning into lipid-rafts (MAM) as well as γ-secretase activity for Aβ generation. These amyloidogenic products have detrimental effects on the cellular lipid homeostasis. Depletion of GSAP maintains a lipid environment with up-regulated phosphatidylethanolamine (PE) and down-regulated ceramide, which improves the bioenergetic capacity of mitochondria. Functionally, we discovered that GSAP deletion restored novel recognitive function in an AD mouse model, and provided evidence that a GSAP SNP is associated elevated GSAP expression correlated with AD.

## FIGURE LEGENDS

**Figure S1:GSAP binding protein and antibody validation(A)** Heatmap showing GSAP and binding protein levels in bait-expressing (GSAP-HA) versus EV (empty vector expression) samples in N2a co-IP and MS analyses. Proteins enriched in GSAP-HA samples are shown; mitochondrial proteins are highlighted in red. **(B)** GO biological process association for GSAP from experimental data and computational prediction (humanbase database, hb.flatironinstitute.org/gene/54103). * is based on previous experimental data. **(C)** GSAP binding protein identified through yeast two hybrid were visualized by the STRING App in Cytoscape. **(D)** Co-IP analysis of GSAP (HA-tagged) interaction with PHB (Flag-tagged) using Flag antibody. **(E)** HA-tagged human GSAP plasmid was transfected into HEK293T cells together with control (C) or GSAP siRNA. 48 hr after transfection, cell lysates were collected and subjected to SDS-PAGE and immunoblot analysis. GSAP antibody from Thermo Scientific (Thermo) or R&D systems (R&D) was used to detect GSAP protein.

**Figure S2:Validation of GSAP gene deletion in mice (A)** Schematic of the gene-targeting strategy to generate GSAP knockout mouse lines (Taconic Farms). Conditional GSAP knockout mice were crossed with CMV-Cre to generate constitutive knockout mouse lines. **(B)** Genomic PCR analysis to distinguish WT (∼325 bp) and GSAP KO (∼500 bp) alleles. **(C)** Quantitative PCR analysis in both WT and GSAP KO mouse hippocampal tissues using primer sets across different GSAP exons. **(D)** Weight was measured for mice used for behavioral studies at 6-months of age. Data represent mean ± s.e.m., unpaired t-test, *p < 0.05, **p < 0.01, N.S. not significant.

**Figure S3:Sn-RNAseq analysis (A)** Distribution profile of nuclei based on cell type (upper panel) or genotype (lower panel). **(B)** Volcano plot showing differentially expressed genes in different clusters in GKO versus WT or J20;GKO versus J20:WT. Only genes with significant expression level change are shown (adjust p-value <0.05; log2(fold change) < -0.3 or > 0.3). Genes with higher expression level in GKO are highlighted in red; genes with lower expression level in GKO are highlighted in blue. **(C)** UMAP plot showing marker gene expression levels from different cell clusters. Color intensity corresponds to gene expression level. **(D)** GO biological process pathway analysis for differentially expressed genes in different cell clusters. **(E)** Module-trait relationship heatmap depicting the correlation between WGCNA gene modules and mouse genotypes. Each cell contains the corresponding correlation (top value) and p-value (bottom value).

**Figure S4:GSAP interacts with lipids and regulates lipid homeostasis (A)** Abbreviation of different lipid subclasses. **(B)** Cell lysates were collected from GSAP-HA transfected cells and overlaid onto lipid-coated membranes. HA antibody was used to detect GSAP protein bound to lipids.

**Figure S5:SNPs affecting GSAP expression in human brain tissues (A)** Human GSAP transcript levels are up-regulated with age in different brain regions. Data were obtained from PsychENCODE. **(B)** Variants located in the GSAP locus identified from European-descent population with significant association with AD (p < 0.05). Data were retrieved from the meta-analysis results from Jansen et al and rank by p-values. β, effect size; BP, base pair; ccREs, candidate cis-regulatory elements; Chr, chromosome; EA, effective allele; NA, not avaliable; SE, stardard error; EAF, effective allele frequency; SNP, single nucleotide polymorphism; TF, transcription factor. **(C)** AD-associated GSAP variants resided in the candidate cis-regulatory elements as reported by the ENCODE Screen database (hg19) listed by genomic coordinates. ccREs, candidate cis-regulatory elements; SNP, single nucleotide polymorphism; TF, transcription factor.

## METHODS

### Mouse Strains

All animal experiments were approved by the Rockefeller University Institutional Animal Care and Use Committee. Mice were maintained in a C57BL/6N genetic background and housed in rooms on a 12 h dark/light cycle interval with food and water available ad libitum. GSAP conditional knockout mice were constructed at Taconic farms by targeting exon 9 to 11 of GSAP. Constitutive GSAP knockout mice were generated by crossing GSAP conditional knockout mice with CMV-Cre mice. J20 mice were purchased from The Jackson Laboratory. Both male and female littermates obtained from in vitro fertilization were used for behavioral tests and single nuclei RNAseq.

### Novel Object Recognition Test

During the habituation phase, mice were placed in an empty open arena for 10 min. 24 h later, mice were placed into the empty arena again for 5 min for the re-habituation phase. Subsequently, two identical objects were fixed to the floor in two corners of the box, and mice were allowed to explore for 10 min. 24 hr later, the familiar object was replaced by a novel object, and mice were allowed to explore for 10 min for the choice phase. Mice interacting with an object for less than 2 sec were removed from analysis. Time spent exploring the objects was recorded during the choice phase. The preference index was quantified as follows: preference index = novel object exploration time/ (novel object exploration time + familiar object exploration time).

### Singe Nuclei RNA Sequencing

Single nuclei RNA sequencing was performed using WT (7-month old; 3 mice), GKO (7-month old; 3 mice), J20;WT (6-month old; 1 mouse), and J20;GKO mice (6-month old; 1 mouse). Single nuclei were isolated based on the published protocol with modifications (Krishnaswami et al., 2016). After dissection, hippocampi were homogenized in a cold Dounce homogenizer after dissection. The homogenate was centrifuged at 1,000g for 8 min at 4°C to pellet nuclei. Nuclei were resuspended in 1 ml 29% iodixanol buffer and centrifuged at 13,500 g for 20 min at 4°C. Supernatant and floating myelin were removed after centrifugation. Nuclei were resuspended in 100 µl nuclei storage buffer and filtered using a 40 µm cell strainer. Nuclei were stained with trypan blue and counted using a hemocytometer. Nuclei (5,000 from each sample) were used for single-nuclei library preparation using the 10X Genomics platform according to the manufacturer’s protocol. Libraries were sequenced on the Novaseq platform. Sample demultiplexing, barcode processing and single-cell counting was performed using the Cell Ranger Single-Cell Software Suite (10x Genomics, version 3.0.2). Cellranger count was used to align samples to the reference genome (mm10). The counting matrix were imported into Seurat package (version 3.0) in R (version 3.6.2) for subsequent analysis. For quality control, nuclei with mitochondrial content >5%, gene number < 200 or gene number > 7500 were removed. After filtering, a total of 32037 individual nuclei across all genotypes were selected for downstream analysis. Data were normalized using a scaling factor of 10,000 by default, and then umi count were normalized using regularized negative binomial regression. Before integration of the 8 samples, 3000 genes were selected by using SelectIntegrationFeatures as the anchor features. The principal component analysis for the integrated dataset were performed using the first 30 principal components and t-SNE analysis was performed with the top 30 PCAs. Clustering was performed using a resolution of 0.8. The raw counting matrix from the Cellranger count were subjected to dimensionality reduction using a ZINB regression model with gene and cell-level covariates. Differential expression of genes (DEGs) between conditions was assessed using the DESeq2. The Excitatory Neurons were selected from the whole single cell dataset according to the cell annotation. The same process was performed on the Excitatory Neuron dataset but with the cluster resolution parameter set as 0.03 in the Seurat package. WGCNA analysis were performed with default parameters on the matrix of raw counts of the Excitatory Neurons and the trait information (genotype). Significant DEGs were used for GO biological pathway analysis using EnrichR (Chen et al., 2013). Meta-enrichment analyses were performed using Metascape. Raw and processed sequencing data reported in this paper are available under GEO accession: GSE157985.

### Cell Culture and Transfection

Mouse N2a neuroblastoma and N2a695 (overexpressing APP695) cells were grown in medium containing 50% DMEM and 50% Opti-MEM, supplemented with 5% FBS and 200 μg/mL G418 (for N2a695) (Life Technologies). HEK293T cells (ATCC; CRL-11268) and IMG (Immortalized adult mouse microglia) cells (Kerafast; EF4001) were grown in DMEM medium containing 10% FBS. SH-SY5Y wild-type and GSAP knockout cells were grown in DMEM-F12 medium containing 10% FBS (Wong et al., 2019). CAD (mouse catecholaminergic neuronal) cells were grown in DMEM-F12 medium containing 8% FBS (Qi et al., 1997). Lipofectamine 2000 and 3000 (Life Technologies) were used for all transient transfections following manufacturer’s instructions. Myc-Flag tagged Arcn1 (RC210778), Phb (RC201229), and Erlin2 (RC221700) were obtained from Origene. Mouse full length GSAP with HA tag (EX-Mm30424-M07), human GSAP plasmids (EX-Z2830-M07), human GSAP-16k with HA tag (a.a. 733 to a.a. 854 subcloned from full length GSAP-HA), Fe65 with Flag tag (EX-M0439-M12), and Fe65 with mCherry tag (EX-Mm20316-M56) were obtained from Genecopoeia. GFP-PP1gamma (gift from Angus Lamond & Laura Trinkle-Mulcahy; #44225) and pSpCas9(BB)-2A-Puro (gift from Feng Zhang; #48139) were obtained from Addgene. Mouse GSAP siRNA, human GSAP siRNA and negative control siRNA were obtained from Dharmacon (On-TARGET plus J-056450-11, LQ-025410-02-0005 and D-001830-02-05).

### SDS-PAGE Immunoblotting and Immunoprecipitation (IP)

Cells were collected and washed with PBS, then lysed with either 3% SDS or RIPA lysis buffer supplemented with protease inhibitor cocktail. Bicinchoninic acid (BCA) assay was used to determine protein concentration. Equal amounts of protein were subjected to SDS-PAGE using either 10-20% Tris-HCl or 4-12% Bis-Tris precast gels. Proteins were transferred onto a PVDF membrane, blocked in 5% non-fat milk for 1 hr at room temperature and incubated with primary antibodies at 4°C overnight. The following primary antibodies were used: Psen1 antibody (1:1,000, MAB5232, EMD Millipore), APP C-terminal antibody (1:4,000, RU369, in house), β-Amyloid antibody 6E10 (1:500, 803001, Biolegend), Phospho-APP (Thr668) antibody (1:1000, 3823S, Cell Signaling Technology), GSAP antibody (1:1000, AF8037, R&D systems), GSAP antibody (1:1000, PA5-21092, ThermoFisher Scientific), PP1β antibody (1:1,000, 07-1217, EMD Millipore), Fe65 antibody (1:1000, ab91650, Abcam), Fe65 antibody (1:1000, sc-19751, Santa Cruz), Phb antibody (1:1000, ab28172, Abcam), Calnexin antibody (1:1000, ab22595, Abcam), HMGCS1 antibody (1:1000, 42201S, Cell Signaling), SREBP2 antibody (1:1000, ab30682, Abcam), Flag M2 antibody (1:1,000, F3165, Sigma), HA antibody (1:1,000, A190-108A, Bethyl), GAPDH antibody (1:500, sc-365062, Santa Cruz), β-tubulin antibody (1:2000, ab6046, Abcam), GFP antibody (1:1000, ab183734, Abcam). Primary antibodies were detected using HRP-linked secondary antibodies together with Western Lightning Plus-ECL (Perkin Elmer). Fiji (ImageJ) was used to quantify band intensity.

For IP experiments, cell pellets were washed with PBS before lysing in IP lysis buffer (50 mM Tris-HCl, 150 mM NaCl, 1% CHAPSO, pH 7.4, supplemented with protease inhibitor cocktail, and PhosStop), for 10 min on ice. Lysates were then centrifuged at 13,000 g for 10 min at 4°C. Prior to IP, supernatants were collected and diluted in IP lysis buffer to reach a CHAPSO final concentration of 0.25%. Primary antibody or IgG control was incubated with lysates overnight at 4°C with tumbling. The next day, 30 μl protein G magnetic beads (ThermoFisher Scientific) were added into samples for 2 h incubation at 4°C. Protein G magnetic beads were collected and washed 4 times with lysis buffer containing 0.25% CHAPSO. Immunoprecipitated proteins were eluted with SDS sample buffer supplemented with reducing reagent. Samples were heated at 70°C for 10 min before subjecting to immunoblot analysis.

### Mass-spectrometry for Binding Protein Identification

HA or GSAP antibody was covalently conjugated to Dynabeads® M-270 Epoxy beads (#14301, Thermo Fisher Scientific) using the antibody coupling kit (#14311D, Thermo Fisher Scientific). Cells cultured in triplicate were lysed in 1% CHAPSO IP lysis buffer (50 mM Tris-HCl, 150 mM NaCl, pH 7.4, supplemented with protease inhibitor cocktail, and PhosStop) and diluted in IP lysis buffer to reach CHAPSO final concentration of 0.25%. HA or GSAP antibody conjugated beads were added into the lysate to tumble for 2 hr at 4°C. Magnetic beads were then collected and washed 3 times with 0.25% CHAPSO IP lysis buffer and 3 times with PBS. The immunoprecipitates were eluted with 8 M urea. Proteins were digested overnight with Endopeptidase Lys-C and trypsin. Peptides were analyzed by nano LC-MS/MS. Data were processed using MaxQuant. Comparing bait versus control samples, the differential enriched protein was labeled as a GSAP binding protein candidate when it had either an average difference > 1.5 or p-value < 0.05. Candidates were subjected to GO biological pathway analysis using DAVID 6.8 (https://david.ncifcrf.gov/). Meta-enrichment analyses were performed using Metascape (Zhou et al., 2019).

### Yeast Two-Hyrid Screening

Yeast two-hybrid screening was performed using the mating strategy with two *Saccharomyces cerevisiae* strains of opposite mating types (strains CG1945 and Y187) as explained elsewhere (Flajolet et al., 2008). The carboxyl terminal of human GSAP cDNA fragment (amino acids from position 497 to 854) was sub-cloned in frame with the GAL4-DNA-BD moiety into a pAS2 vector as the bait following standard procedures. The bait construct expression was evaluated prior screening by western blotting analysis (anti-GAL4 domain antibody) after transfection of the bait plasmids in yeast. Toxicity and auto-activity levels of the bait were also evaluated. A commercial cDNA human brain library (sub-cloned into pACTII) was used and served as the prey. Plasmids of positive clones growing on selective medium were rescued and submitted to DNA sequencing for clone identification using NCBI-BLAST.

### ELISA for Aβ

Aβ quantification was performed as described in our previous publication (Bettayeb et al., 2016a). Briefly, wild-type and Fe65KO CAD cells were transiently transfected with APP constructs. Media were replaced 6 h before collecting supernatants. Conditioned media from CAD cells were then diluted in buffer for Aβ measurement following the manufacturer’s instructions (Thermo Fisher Scientific). Aβ levels were normalized to total protein levels.

### MAM Subcellular Fractionation

MAM subcellular fractionation was performed as previously described (Lewis et al., 2016). Briefly, cells were homogenized in a sucrose buffer (0.25 M sucrose) using a Teflon glass homogenizer. Homogenates were centrifuged at 600 g for 5 min. Pellets were resuspended in isolation medium (5 mM HEPES, pH 7.4, 250 mM mannitol, 0.5 mM EGTA) and centrifuged at 10,300 g for 20 min. Supernatants were centrifuged at 100,000 g for 1 h to separate the microsome and cytosol fractions. Pellets were resuspended in isolation medium and layered on top of a Percoll medium and centrifuged at 95,000 g for 30 min. MAM and mitochondria fractions were collected from different layers after centrifugation. The MAM fraction was centrifuged at 100,000 g for 1 h to obtain MAM.

### Immunofluorescence Microscopy

For immunofluorescence microscopy, cells were fixed in 4% paraformaldehyde for 10 min, mounted using Vectashield mounting medium with DAPI (Vector Laboratories, Inc.) and analyzed using a Zeiss LSM710 Fluorescence Microscope. For live cell imaging, super-resolution images were acquired using a Zeiss LSM 800 confocal microscope equipped with Airyscan module (Zeiss). Fluorescence was collected with ×40 objective lens. Vesicle trafficking velocity and diffusion coefficient were calculated by MATLAB.

### Electron Microscopy

Cells grown on ACLAR film were fixed with 4% formaldehyde and 2% glutaraldehyde in 0.1M sodium cacodylate buffer (pH 7.4). Subsequently, cells were washed in the buffer, post-fixed with 1% osmium tetra-oxide for 1 h, stained en bloc with 1% uranyl acetate for 30 min, dehydrated by a graded series of ethanol, infiltrated with a resin (Eponate12, Electron Microscope Sciences) and embedded with the resin. After polymerization at 60°C for 48 h, ultra-thin sections were cut, underwent post-staining with 2% uranyl acetate and 1% lead citrate and were examined under a JEOL 1400Plus transmission electron microscope.

### Generation of Fe65 Knockout CAD line

Guide RNA sequence (5′-acggattccgatctaccggc-3′) targeting the mouse Fe65 gene was cloned into the pSpCas9(BB)-2A-Puro vector, a gift from Feng Zhang (Addgene plasmid #48139). The plasmid was transfected into CAD cells, which underwent 1 μg/ml puromycin selection 48 h after transfection. Cells were seeded in clonal limiting dilution in 96 well plates. Fe65KO cells were screened and validated by immunoblot analysis and Sanger sequencing.

### Lipidomics Analysis

Lipid extracts were prepared using a modified Bligh and Dyer method (Bligh and Dyer, 1959). Extracts were spiked with appropriate internal standards and analyzed by LC/MS system as described (Chan et al., 2012). Briefly, glycerophospholipids and sphingolipids were separated with normal-phase high performance liquid chromatography (HPLC), while sterols and glycerolipids were separated with reverse-phase HPLC using an isocratic mobile phase. Individual lipid species were quantified by referencing to spiked internal standards. The nomenclature abbreviations are listed in Figure S5B.

### Lipid Overlay Assay

A nitrocellulose membrane spotted with the indicated lipids (Echelon Biosciences) was blocked in 3% BSA in PBST (0.1% Tween 20) at 4°C overnight. HEK293T cells transiently expressing HA-GSAP were lysed in 0.5% Triton lysis buffer (50 mM Tris-HCl, 150mM NaCl, pH 7.4). Cell lysate (200 µg) was diluted in 3% BSA in PBST for 1 h incubation with the membrane at room temperature. After washing, GSAP association with lipids was detected using HA antibody and HRP-conjugated secondary antibody.

### Cellular Respiration Analysis

Oxygen consumption reflecting mitochondrial activity was measured by XF mito stress kit according to the manufacturer’s protocol. All the measurements were performed using the Agilent Seahorse XFe96 analyzer from the HTSRC. WT and GSAP KO SHSY-5Y cells were seeded at 20,000 per well in the 96-well plate one day before the measurement. Oxygen consumption rate was measured after sequential addition of 1 μM oligomycin, 1 μM FCCP and 0.5 μM rotenone/antimycin. The results were analyzed using the Wave software (Agilent) and normalized by the cell number, which was measured by the ImageXpress-micro system.

### Identification and Annotation of AD-Associated Genetic Variants in GSAP Locus

AD association of GSAP variants (GRCh37, chr9: 76890110-77095630) was obtained from the recent published AD GWAS summary statistics (Jansen et al., 2019). GSAP variants with P < 0.05 were retained as variants exerting AD association. The obtained AD-associated GSAP variants were subjected to the SCREEN hg19 database (https://screen-v10.wenglab.org/gwasApp/?assembly=hg19; (Consortium et al., 2020)) for annotating variants that may reside in the candidate cis-regulatory elements (cCREs). IGV (Integrative Genomics Viewer, version 2.8.7) was used to visualize the epigenic events in rs74338692-assocated genomic regions. Specifically, the following datasets were analyzed in the study:

> Cell-type agnostic ccRE (ENCODE ID: ENCFF788SJC) Monocytes:
>
> DNase-seq (ENCODE ID: ENCFF398USK)
>
> H3K27ac (ENCODE ID: ENCFF931PZJ)
>
> H3K4me1 (ENCODE ID: ENCFF731YSQ)
>
> Bipolar neurons:
>
> DNase-seq (ENCODE ID: ENCFF106BSM)
>
> H3K27ac (ENCODE ID: ENCFF967OEW)
>
> Transcription factor binding events:
>
> FOSL2 (ENCODE ID: ENCFF321KVH)

### Statistical Analysis and Data Visualization for GSAP Variants

Linkage disequilibrium and haplotype analysis for six GSAP variants residing in the cCREs were conducted using 1000 Genomes Project Phase 3 whole-genome sequencing data of European Super Population (EUR, n = 503). In brief, genotypes for those 6 SNPs stored in VCF files were extracted and subjected to PLINK (version v1.90b6.12; (Purcell et al., 2007)) for analysis. The ped file obtained from PLINK analysis was subsequently subjected to Haploview (version 4.2; (Barrett et al., 2005)) for linkage disequilibrium and haplotype analysis and visualization. For genotype-expression association analysis, the whole-genome sequencing genotype information obtained from the GTEx (phs000424.v8.p2) were further subjected to BEAGLE (version r1399) for haplotype phasing (nthreads=24, phase-its=50, impute-its=30). The haplotypes constructed by 6 variants were obtained by R programming analysis of phased genotypes. Association analysis was conducted between GSAP variant or haplotype dosage, and GSAP transcript levels in 13 brain regions recorded in GTEx database by robust regression analysis (R robustbase packages). Meta-analysis was further carried out by summarizing results at tissue levels using METASOFT (version 2.0.1). GraphPad Prism (version 8.0.1) was used to generate bar and dot plots.

### Statistical Analysis

Statistical analysis was performed by using GraphPad Prism. Results are presented as mean ± s.e.m or mean ± s.e as indicated. Matlab was used for live cell imaging data analysis. Two-tailed unpaired Student’s t-test was used, except for sn-RNAseq analysis. p < 0.05 was considered significant (*p < 0.05, **p < 0.01, ***p < 0.001 and ****p < 0.0001). For the animal behavior study, mice from the same litter were randomnized into groups and was the experiment was performed blined.

## ACKNOWLEDGEMENTS

We would like to thank Drs. C. Zhao, and H. Duan in the Genomics Resource Center; Dr. H. Molina from the Proteomics Resource Center; R. Norinsky, R. Cubias and J. Torres in the Transgenic Services Resource Center; Drs. K. Uryu and N. Soplop in the Electron Microscope Resource Center; Drs L. Ramos-Espiritu and J.F. Glickman in the high throughput and spectroscopy resource center (HTSRC) for their excellent technical service. Authors also acknowledge the MSK Cancer Center Support Grant/Core Grant (Grant P30 CA008748) and MSK Molecular Cytology Core Facility/Core Grant (P30 CA 008748). We thank Dr. E. Area-Gomez for performing the lipidomic analysis; Dr. X. Fan for the help on figure plotting. This work was supported by Fisher Center for Alzheimer’s Research (P.G. and M.F.), NIH/NIA AG061350 (Y.M.L. and M.F.) and JPB Foundation (Y.M.L., P.G. and M.F.). Research in the laboratory of A.C.N. is supported by the National Institutes of Health (NIH) (AG047270, AG062306, AG066508, DA018343) and the State of Connecticut Department of Mental Health and Addiction Services. This paper is dedicated to our beloved mentor, the late Dr. Paul Greengard.

## AUTHOR CONTRIBUTIONS

P.X., J.C.C., Y.M.L., and P.G. conceived the study. P.X. analyzed data from proteomics, sn-RNAseq, lipidomics, Cellular respiration, animal behavioral experiments and performed biochemical, immunofluorescence and constructed Fe65 CRISPR-Cas9 knockout cells. J.C.C. performed live cell imaging experiments and analyzed data. X.Z. performed the human genetic analysis. W.W. performed bioinformatic analysis for sn-RNAseq and visualized data with P.X.. M.B. generated the mouse cohort and performed behavioral tests. E.W. generated the GSAP CRISPR-Cas9 knockout cells. K.B. performed the Y2H experiment. P.X., L.J. and T.H. analyzed the EM data. T.H. and A.C.N.edited the manuscript. W.L. constructed the GSAP knockout mice. P.X., H.X., A.C.N., M.F., N.I., Y.M.L., and P.G. designed the project and supervised experiments. P.X. summarized findings in illustrations. P.X., and Y.M.L. wrote the manuscript.

## DECLARATION OF INTERESTS

Y.M.L. is a co-inventor of the intellectual property (assay for γ-secretase activity and screening method for γ-secretase inhibitors) owned by MSKCC and licensed to Jiangsu Continental Medical Development.

