## Supplementary material for "GSAP regulates mitochondrial function through the Mitochondria-associated ER membrane in the pathogenesis of Alzheimer’s disease": Fig. S1

A

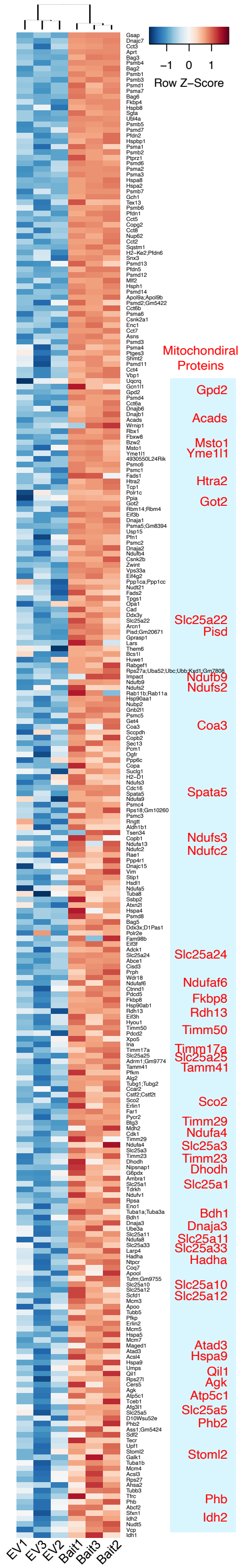


B

| NAME | IDENTIFIER | CONFIDENCE | SOURCE |
|---|---|---|---|
| regulation of cellular amide metabolic process | GO:0034248 | 1 | Experimentally associated* |
| positive regulation of cellular amide metabolic process | GO:0034250 | 1 | |
| peptide metabolic process | GO:0006518 | 1 | |
| amyloid-beta formation | GO:0034205 | 1 | |
| amyloid precursor protein catabolic process | GO:0042987 | 1 | |
| amyloid-beta metabolic process | GO:0050435 | 1 | |
| amyloid precursor protein metabolic process | GO:0042982 | 1 | |
| regulation of amyloid-beta formation | GO:1902003 | 1 | |
| regulation of amyloid precursor protein catabolic process | GO:1902991 | 1 | |
| positive regulation of amyloid-beta formation | GO:1902004 | 1 | |
| positive regulation of amyloid precursor protein catabolic process | GO:1902993 | 1 | |
| regulation of DNA binding transcription factor activity | GO:0051090 | 0.12757446 | Computationally predicted |
| regulation of immune response | GO:0050776 | 0.123980903 | |
| positive regulation of MAPK cascade | GO:0043410 | 0.11471183 | |
| regulation of protein serine/threonine kinase activity | GO:0071900 | 0.110187344 | |
| positive regulation of protein transport | GO:0051222 | 0.106151562 | |
| positive regulation of immune response | GO:0050778 | 0.104747482 | |
| positive regulation of protein kinase activity | GO:0045860 | 0.10304935 | |
| positive regulation of cytokine production | GO:0001819 | 0.101247148 | |

C

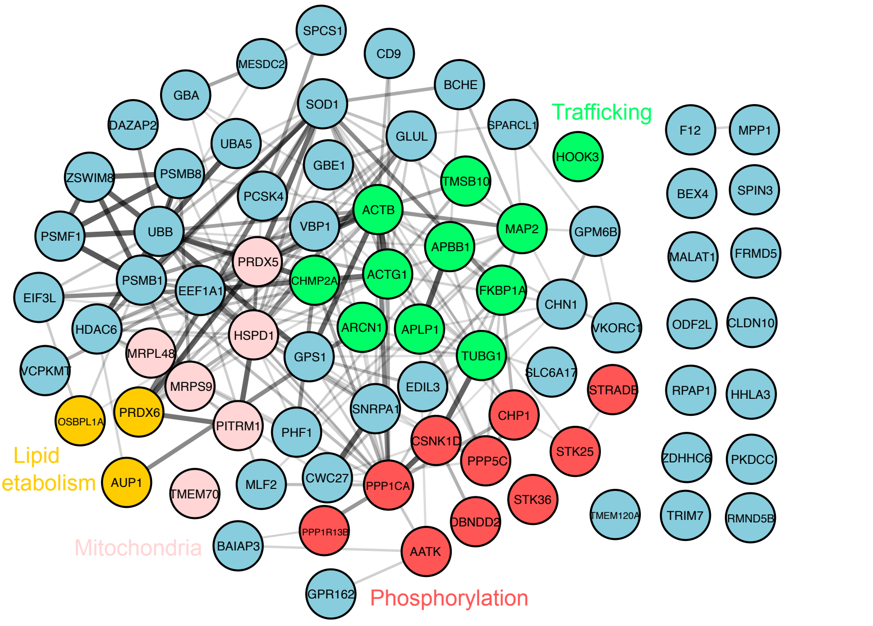


D

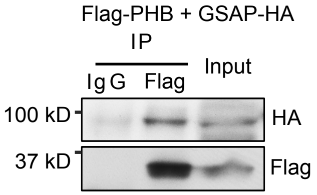


E

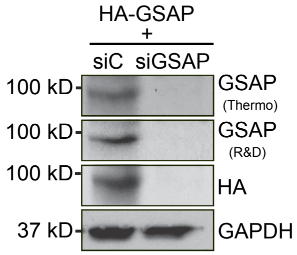


Fig. S1
