## Supplementary figures and images for "GSAP regulates mitochondrial function through the Mitochondria-associated ER membrane in the pathogenesis of Alzheimer’s disease"

### Fig. S2

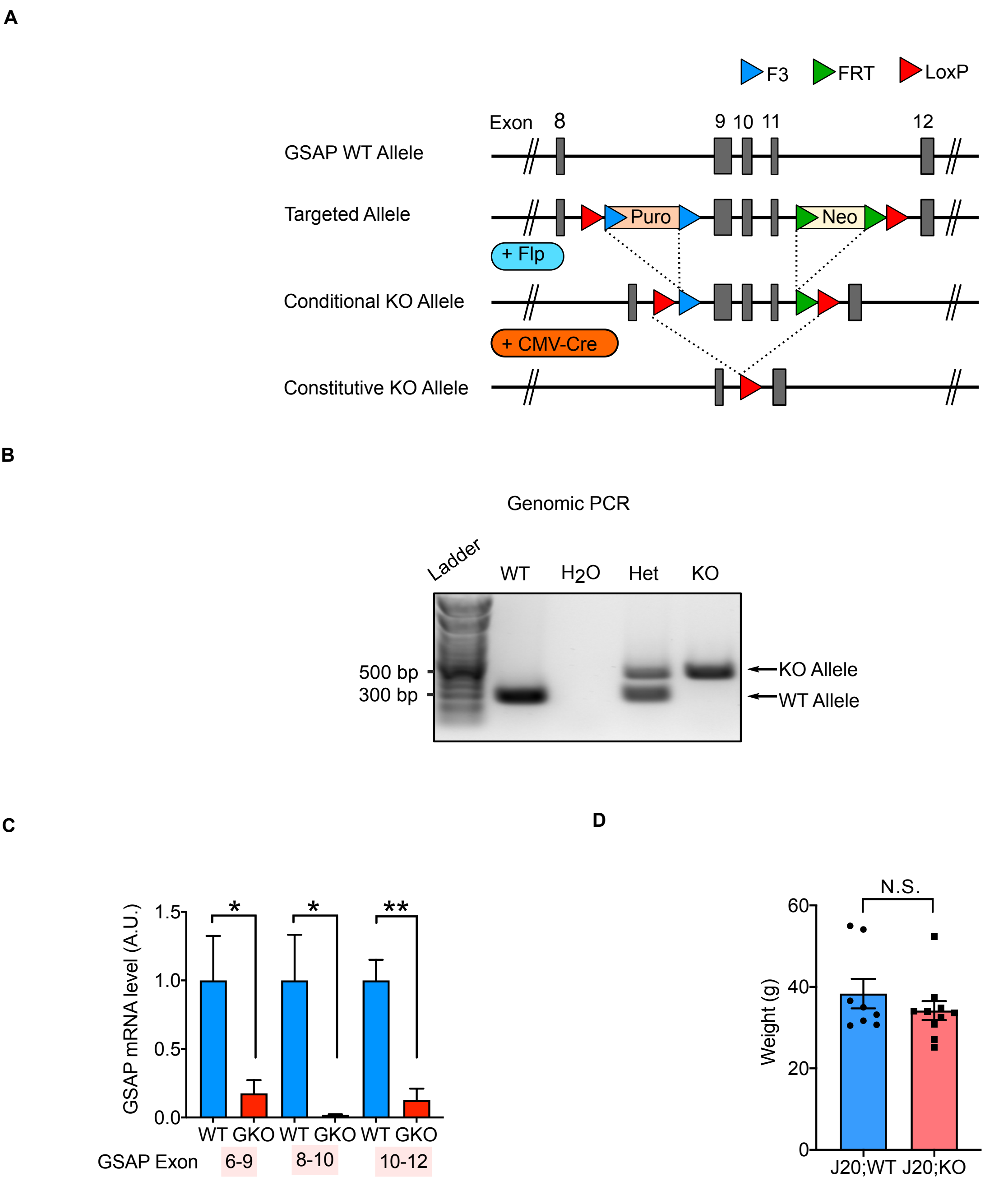

### Fig. S3

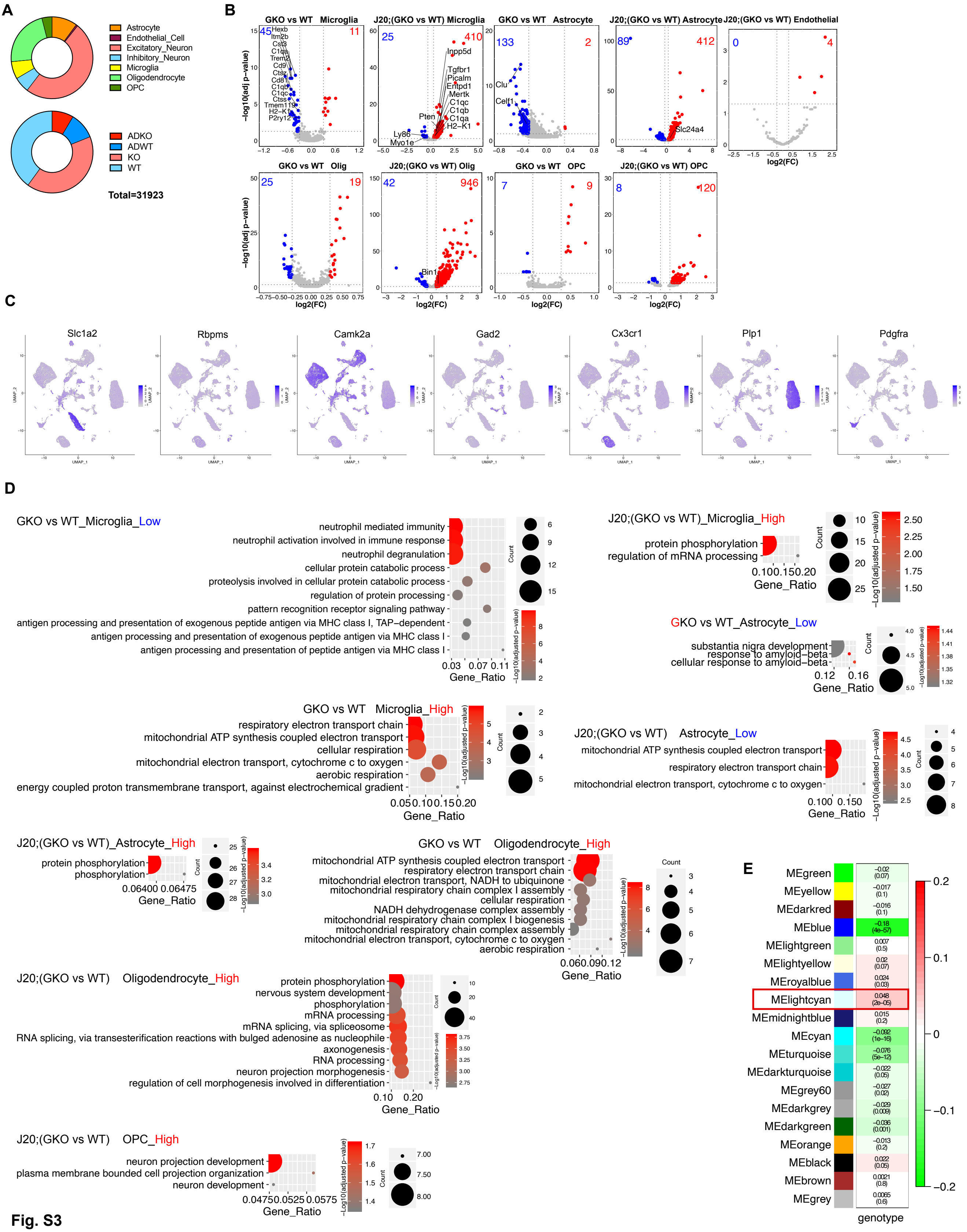
