## Supplementary material for "GSAP regulates mitochondrial function through the Mitochondria-associated ER membrane in the pathogenesis of Alzheimer’s disease": Fig. S4

**A**

| | | | |
|---|---|---|---|
| **FC** | Free Cholesterol | **PCe** | Ether phosphatidylcholine |
| **CE** | Cholesterol Ester | **PE** | Phosphatidylethanolamine |
| **AC** | Acyl Carnitine | **PEp** | Plasmalogen phosphatidylethanolamine |
| **MG** | Monoacylglycerol | **PS** | Phosphatidylserine |
| **DG** | Diacylglycerol | **PI** | Phosphatidylinositol |
| **TG** | Triacylglycerol | **PG** | Phosphatidylglycerol |
| **dhCer** | Dihydroceramide | **BMP** | Bis(monoacylglycero)phosphate |
| **Cer** | Ceramide | **AcylPG** | Acyl Phosphatidylglycerol |
| **SM** | Sphingomyelin | **LPC** | Lysophosphatidylcholine |
| **dhSM** | Dihydrosphingomyelin | **LPCe** | Ether lysophosphatidylcholine |
| **Sulf** | Sulfatide | **LPE** | Lysophosphatidylethanolamine |
| **MHCer** | Monohexosylceramide | **LPEp** | Plasmogen Lysophosphatidylethanolamine |
| **LacCer** | Lactosylceramide | **LPI** | Lysophosphatidylinositol |
| **GM3** | Monosialodihexosylganglioside | **LPS** | Lysophosphatidylserine |
| **GB3** | Globotriaosylceramide | **NAPE** | N-Acyl Phosphatidylethanolamine |
| **PA** | Phosphatidic acid | **NAPS** | N-Acyl Phosphatidylserine |
| **PC** | Phosphatylcholine | **NSer** | N-Acyl Serine |

**B**

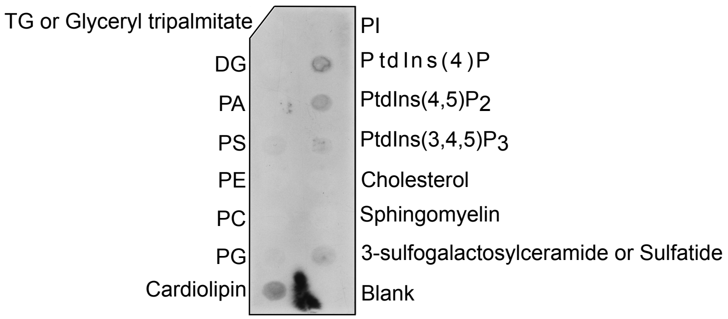


**Fig. S4**
