## Supplementary material for "GSAP regulates mitochondrial function through the Mitochondria-associated ER membrane in the pathogenesis of Alzheimer’s disease": Fig. S5

**A**

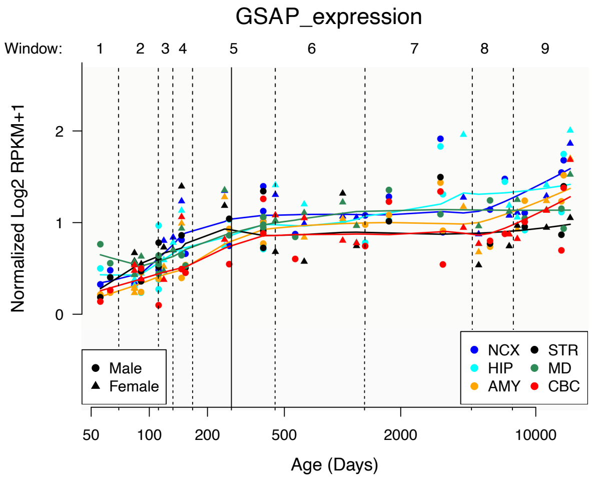


**B**

| SNP | Chr:BP (GRCh37) | EA | Z | β | SE | p | EAF | TF binding | ccREs_ID |
|---|---|---|---|---|---|---|---|---|---|
| rs10274222 | 7:76958730 | A | 2.599 | 0.01 | 0.004 | 9.35E−03 | 0.092 | No | EH37E1276447 |
| rs6965851 | 7:76949533 | A | 2.586 | 0.01 | 0.004 | 9.70E−03 | 0.096 | Yes | EH37E0910175 |
| rs149313725 | 7:76926211 | A | −2.466 | −0.022 | 0.009 | 1.37E−02 | 0.018 | No | NA |
| rs6950444 | 7:76978096 | A | 2.431 | 0.011 | 0.004 | 1.51E−02 | 0.066 | Yes | EH37E0910197 |
| rs62476214 | 7:77062290 | C | −2.427 | −0.011 | 0.004 | 1.52E−02 | 0.067 | No | NA |
| rs62476215 | 7:77065751 | T | −2.281 | −0.010 | 0.004 | 2.26E−02 | 0.067 | No | NA |
| rs28378429 | 7:77064886 | A | −2.268 | −0.010 | 0.004 | 2.33E−02 | 0.069 | No | NA |
| rs62476216 | 7:77067081 | A | −2.256 | −0.010 | 0.004 | 2.40E−02 | 0.067 | No | NA |
| rs75611871 | 7:76907606 | A | −2.203 | −0.014 | 0.006 | 2.76E−02 | 0.031 | No | NA |
| rs116993213 | 7:76967512 | G | 2.19 | 0.009 | 0.004 | 2.85E−02 | 0.078 | No | EH37E0910183 |
| rs147072075 | 7:76929790 | G | −2.178 | −0.018 | 0.008 | 2.94E−02 | 0.018 | No | NA |
| rs770596596 | 7:77032560 | C | 2.097 | 0.236 | 0.113 | 3.60E−02 | 0 | No | NA |
| rs74338692 | 7:76946200 | T | 2.046 | 0.016 | 0.008 | 4.07E−02 | 0.018 | Yes | EH37E0910174 |
| rs6465530 | 7:76913475 | A | 1.985 | 0.008 | 0.004 | 4.72E−02 | 0.088 | Yes | EH37E1276433 |
| rs28674557 | 7:76959908 | T | −1.973 | −0.030 | 0.015 | 4.85E−02 | 0.15 | Yes | NA |
| rs60834657 | 7:76996152 | G | −1.963 | −0.006 | 0.003 | 4.96E−02 | 0.158 | No | NA |

**C**

| SNP | β (AD) | EAF | TF binding | ccREs_ID | Z score | | | |
|---|---|---|---|---|---|---|---|---|
| | | | | | DNase | H3K4me3 | H3K27ac | CTCF |
| rs6465530 | 0.008 | 0.088 | Yes | EH37E1276433 | 3.19 | 1.69 | 3.74 | 2.58 |
| rs74338692 | 0.016 | 0.018 | Yes | EH37E0910174 | 2.69 | 2.71 | 4.01 | 1.31 |
| rs6965851 | 0.01 | 0.096 | Yes | EH37E0910175 | 3 | 1.46 | 2.52 | 1.34 |
| rs10274222 | 0.01 | 0.092 | No | EH37E1276447 | 2.72 | 3.42 | 3.83 | 1.57 |
| rs116993213 | 0.009 | 0.078 | No | EH37E0910183 | 2.39 | 2.8 | 3.08 | 1.2 |
| rs6950444 | 0.011 | 0.066 | Yes | EH37E0910197 | 2.73 | 1.18 | 1.71 | 1.34 |

**Fig. S5**
